# Genome-wide profiling of highly similar paralogous genes using HiFi sequencing

**DOI:** 10.1101/2024.04.19.590294

**Authors:** Xiao Chen, Daniel Baker, Egor Dolzhenko, Joseph M Devaney, Jessica Noya, April S Berlyoung, Rhonda Brandon, Kathleen S Hruska, Lucas Lochovsky, Paul Kruszka, Scott Newman, Emily Farrow, Isabelle Thiffault, Tomi Pastinen, Dalia Kasperaviciute, Christian Gilissen, Lisenka Vissers, Alexander Hoischen, Seth Berger, Eric Vilain, Emmanuèle Délot, UCI Genomics Research to Elucidate the Genetics of Rare Diseases (UCI GREGoR) Consortium, Michael A Eberle

**Affiliations:** PacBio, Menlo Park, CA, USA; GeneDx, Gaithersburg, MD, USA; Genomic Medicine Center, Children’s Mercy Kansas City, Kansas City, MO, USA; UMKC School of Medicine, University of Missouri Kansas City, Kansas City, MO, USA; Department of Pediatrics, Children’s Mercy Kansas City, Kansas City, MO, USA; Department of Pathology and Laboratory Medicine, Children’s Mercy Kansas City, Kansas City, MO, USA; Genomics England Ltd., London, UK; Department of Human Genetics, Radboud University Medical Center, Nijmegen, The Netherlands; Research Institute for Medical Innovation, Radboud University Medical Center, Nijmegen, The Netherlands; Radboud Center for Infectious Diseases (RCI), Department of Internal Medicine, Radboud University Medical Center, Nijmegen, The Netherlands; Radboud Expertise Center for Immunodeficiency and Autoinflammation and Radboud Center for Infectious Disease (RCI), Radboud University Medical Center, Nijmegen, The Netherlands; Center for Genetics Medicine Research, Children’s National Hospital, Washington, DC, USA; Institute for Clinical and Translational Science, University of California, Irvine, CA, USA

## Abstract

Variant calling is hindered in segmental duplications by sequence homology. We developed Paraphase, a HiFi-based informatics method that resolves highly similar genes by phasing all haplotypes of a gene family. We applied Paraphase to 160 long (>10 kb) segmental duplication regions across the human genome with high (>99%) sequence similarity, encoding 316 genes. Analysis across five ancestral populations revealed highly variable copy numbers of these regions. We identified 23 families with exceptionally low within-family diversity, where extensive gene conversion and unequal-crossing over have resulted in highly similar gene copies. Furthermore, our analysis of 36 trios identified 7 *de novo* SNVs and 4 *de novo* gene conversion events, 2 of which are non-allelic. Finally, we summarized extensive genetic diversity in 9 medically relevant genes previously considered challenging to genotype. Paraphase provides a framework for resolving gene paralogs, enabling accurate testing in medically relevant genes and population-wide studies of previously inaccessible genes.

## Introduction

Population-wide whole-genome sequencing (WGS) studies based on short reads have enabled comprehensive characterization of variants, particularly small variants, in ∼90% of the human genome (The 1000 Genomes Project Consortium 2015; Bycroft et al. 2018; Karczewski et al. 2020). However, there exist difficult regions and variant classes that remain largely inaccessible to short reads (Mandelker et al. 2016; Ebbert et al. 2019). A large portion of these difficult regions occur within segmental duplications (SDs) (Bailey et al. 2001; Vollger et al. 2022), where high sequence similarity between copies of SDs results in ambiguous mapping of short reads. In addition to difficulty mapping reads within SDs, high sequence similarity promotes unequal crossing over, resulting in hotspots for copy number variants (CNVs), as well as high rates of gene conversion (Chen et al. 2007). These high rates of gene conversion promote sequence exchange between SDs further increasing the errors in read alignment. While short-read based computational methods have been developed to improve the genotyping capability and diagnostic yield in segmental duplications (Handsaker et al. 2015; Ebbert et al. 2019; Prodanov and Bansal 2022; Steyaert et al. 2023), comprehensive variant calling in these regions remains a challenge, and SDs have not been studied at the population level by the current high throughput technologies.

Many medically relevant genes fall into SDs where traditional alignment-based analysis has not been demonstrated to reliably detect the full diversity of these regions. For example, spinal muscular atrophy is caused by variants in the *SMN1* gene, which has a highly similar paralog *SMN2* (Lunn and Wang 2008). Another disease, 21-Hydroxylase-Deficient Congenital Adrenal Hyperplasia (21-OHD CAH), is caused by variants in the *CYP21A2* gene (Merke and Auchus 2020), which resides in a 30-kb tandem repeat called the RCCX module and has a pseudogene *CYP21A1P*. Variants in the *OPN1LW*/*OPN1MW* gene cluster, which contains 1-5 copies of *OPN1LW* or its paralog *OPN1MW*, cause color vision deficiencies (Neitz and Neitz 2011, 2021). To date, these medically important SD-encoded genes are studied with multi-step analyses including a combination of low or medium-throughput assays such as MLPA, amplicon sequencing, or long-range PCR followed by Sanger sequencing to detect copy number (CN) changes or individual variants (Pignatelli et al. 2019; Haer-Wigman et al. 2022). These tests are sometimes limited to a few known variants and may be prone to false negatives if the patient has a pathogenic variant that is not part of the test. There remains a need to fully characterize these genes both for research and clinical testing.

Recently, researchers have begun to study SDs using long-read sequencing. High quality phased assemblies have been generated for a number of samples (Wang et al. 2022; Liao et al. 2023; Gao et al. 2023) using PacBio HiFi and Oxford Nanopore Technologies (ONT) long reads, revealing the sequences of SDs and providing biological and evolutionary insights (Vollger et al. 2022, 2023). However, SDs with multiple copies of highly similar regions are prone to assembly errors especially in regions of extended sequence homology (Vollger et al. 2023). Alternatively, we developed a phasing approach, Paraphase, that identifies haplotypes of genes and their paralogs and demonstrated the ability to accurately resolve the highly similar *SMN1*/*SMN2* region (Chen et al. 2023). That study was limited to one difficult region, leaving a need for a genome-wide demonstration.

Here we extended Paraphase to analyze 316 paralogous genes that fall into 160 groups of SD regions across the genome, including many medically relevant genes that were traditionally considered challenging to genotype. Applying Paraphase to 259 individuals from five ancestral populations, we showed the genetic diversity of these regions across populations in copy number (CN) and sequence variation. We note that some of these regions show exceptionally low diversity between genes and paralogs, signaling high rates of gene conversion. Finally, we studied the Paraphase derived haplotypes for these gene families in 36 parent-offspring trios and identified 11 *de novo* events, among which 7 are *de novo* single nucleotide variants (SNVs) and 4 are consistent with *de novo* gene conversion events.

## Results

### Profiling 160 gene-coding homologous regions with Paraphase

Paraphase resolves highly similar genes by realigning HiFi reads to one, most relevant, gene chosen to represent all copies of the gene and its paralogs. We call this gene the archetype gene. For example, to study *SMN1* and *SMN2*, we realign all of the reads that are aligned to either *SMN1* or *SMN2* to just *SMN1* because that is the fully functional copy. The aligned reads are then phased into haplotypes for variant calling (Figure 1A). For this study, we identified 160 homologous regions >10kb in length with >99% sequence similarity that were found between two and four times in GRCh38 (Table S1, also see Methods). These homologous regions encode 316 genes in total (excluding pseudogenes). In this paper, the term “gene family” is used to describe a set of genes that are highly similar in sequence and are analyzed by Paraphase as a group.

**Figure 1.**
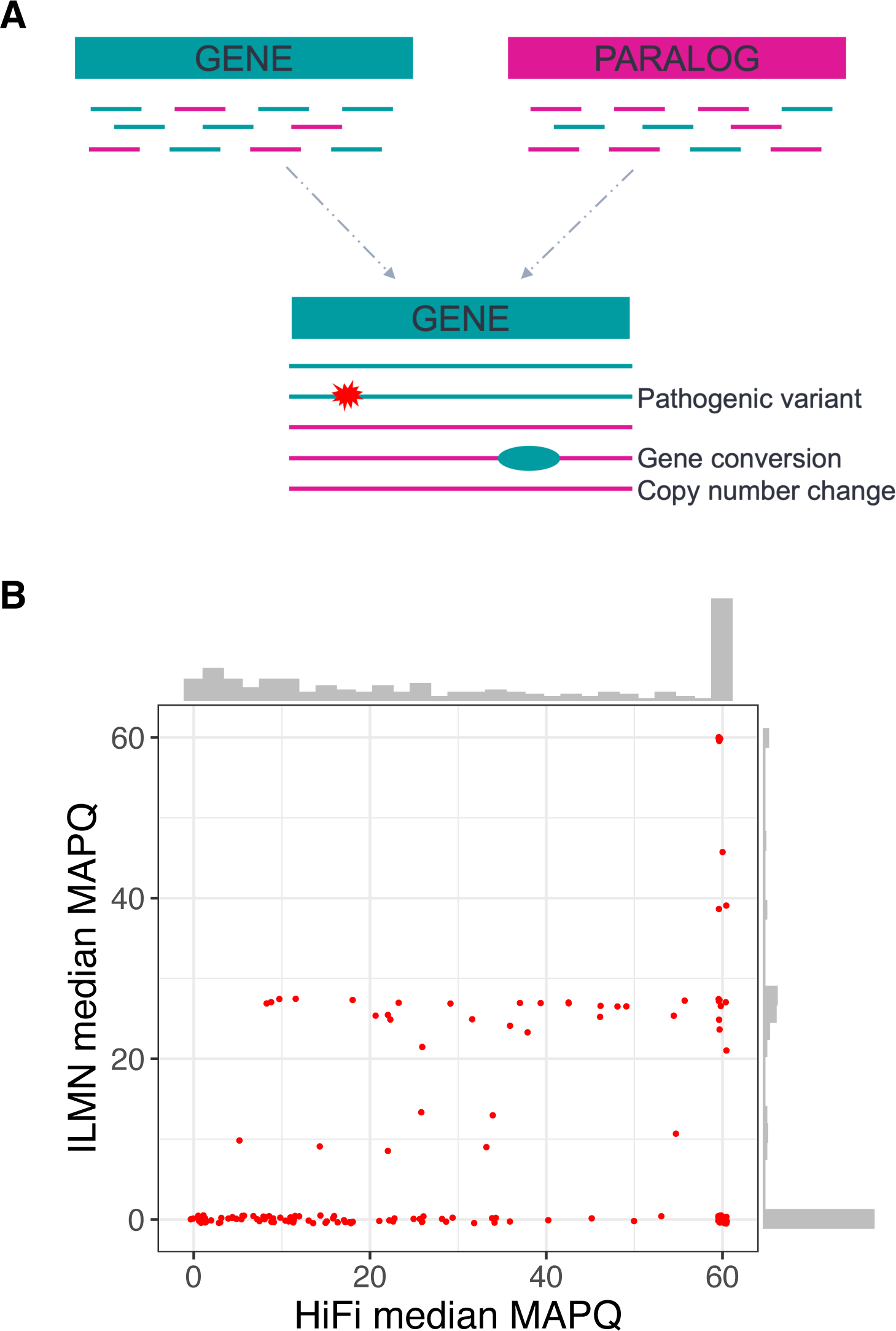
Paraphase design and the regions it analyzes. **A.** Paraphase extracts reads that align to gene families, realigns to the archetype gene, and phases reads into haplotypes. Variant calling is performed on each haplotype. **B**. Comparison of median MAPQ between HiFi and Illumina WGS data in 160 groups of homologous regions analyzed by Paraphase, highlighting mapping difficulty in these challenging regions for both short and long reads.

Among the 160 gene families (Table S1), 149 have gene family members located on the same chromosome, with 16 in tandem (less than 10kb apart). To quantify the impact of sequence homology on read alignments, we examined the mapping qualities (MAPQs) in these regions in both short-read and long-read sequence data (Figure 1B). For short-read data, MAPQs are extremely low (76.4% have a median MAPQ<=20, and 98.8% have genomic segments with MAPQ<=20), indicating the difficulty of mapping short reads to these regions. Even for long-read data, 44.1% of the regions have a median MAPQ<=20 and 75.2% have genomic segments with MAPQ<=20. There are 25 (15.6%) gene families where the median MAPQ is 60 and there are no bases with MAPQ<=20 in the long-read data. These are either regions where the homology extends less than the HiFi read length of ∼15-20 kb or regions included in Paraphase for fusion calling between less similar paralogs (see Methods). Paraphase analysis is still needed in these high MAPQ regions because: 1) reads with high MAPQ can be misaligned due to reference genome artifacts, common CNVs and high rates of gene conversion, 2) gene fusions are hard to detect because split alignments are unlikely to happen in regions of homology and 3) lower MAPQs are expected in data with shorter read length, such as in HiFi hybrid capture data.

### Validation of Paraphase calls

We first validated Paraphase variant calls in 8 medically relevant genes in 21 disease or carrier samples using orthogonal methods such as MLPA and Sanger sequencing (Table 1 and Table S2, also see Methods). For this validation, Paraphase correctly identified all 30 of the clinical variants in these samples.

**Table 1.**
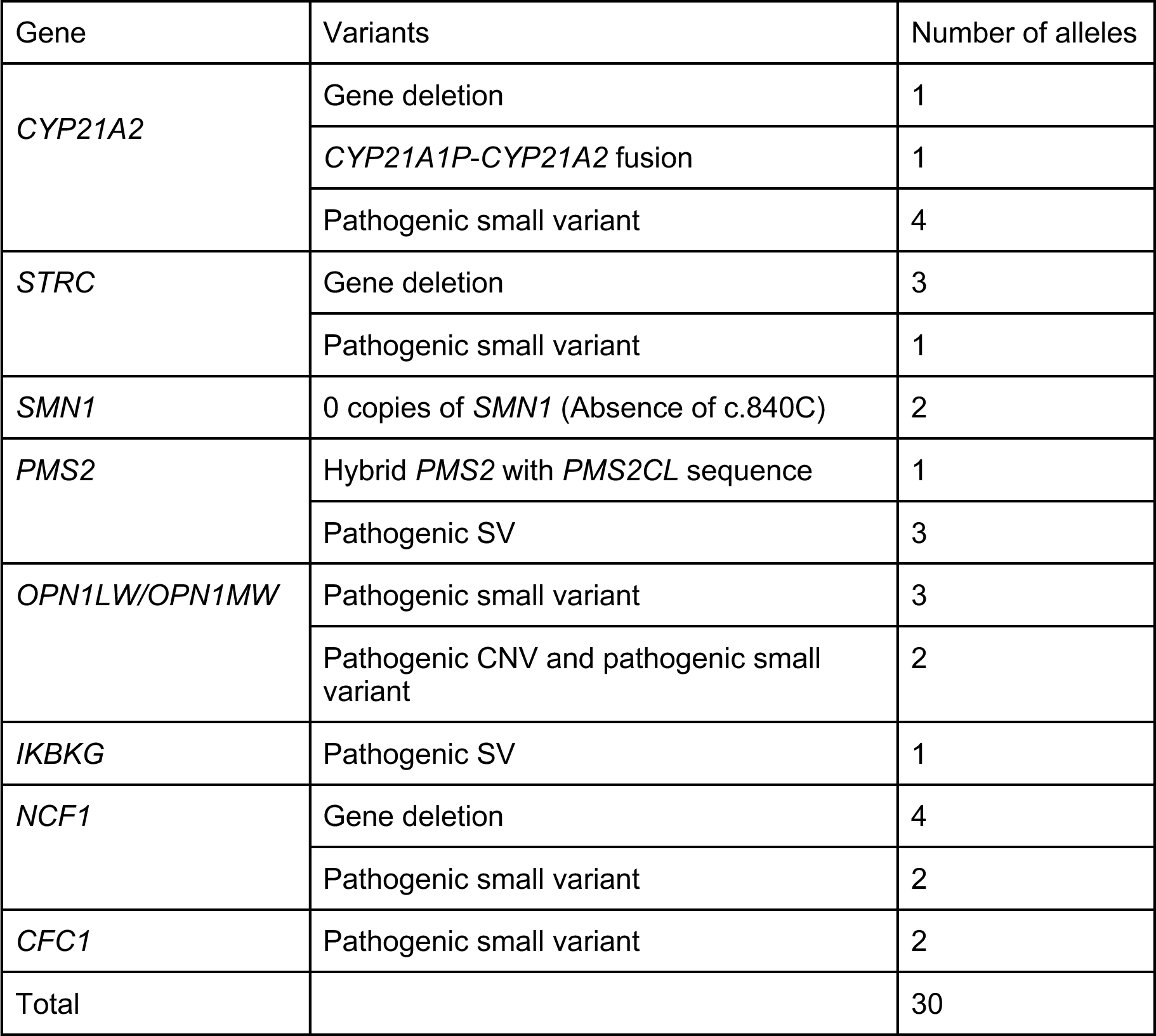
Paraphase calls in medically relevant genes validated by orthogonal methods.

| Gene | Variants | Number of alleles |
| --- | --- | --- |
| *CYP21A2* | Gene deletion | 1 |
|  | *CYP21A1P-CYP21A2* fusion | 1 |
|  | Pathogenic small variant | 4 |
| *STRC* | Gene deletion | 3 |
|  | Pathogenic small variant | 1 |
| *SMN1* | 0 copies of *SMN1* (Absence of c.840C) | 2 |
| *PMS2* | Hybrid *PMS2* with *PMS2CL* sequence | 1 |
|  | Pathogenic SV | 3 |
| *OPN1LW/OPN1MW* | Pathogenic small variant | 3 |
|  | Pathogenic CNV and pathogenic small variant | 2 |
| *IKBKG* | Pathogenic SV | 1 |
| *NCF1* | Gene deletion | 4 |
|  | Pathogenic small variant | 2 |
| *CFC1* | Pathogenic small variant | 2 |
| Total |  | 30 |

We also examined haplotypes called by Paraphase in 36 trios. Among 14,734 full-length haplotypes called in the probands (also requiring full-length haplotypes called in the two parents of each trio), 14,679 (99.6%) agreed exactly with one of the haplotypes observed in the parents. Upon examining the 55 inconsistent haplotypes, 43 (0.29%) are Paraphase errors (switch errors or extra or missing haplotypes), and the remaining 12 (0.081%) are true recombination or *de novo* events (See “Identification of *de novo* mutations and gene conversion” section).

### CN variability of gene families

We calculated the distribution of the total CN (defined by the number of unique haplotypes, adjusted by depth) of each gene family in 259 unrelated individuals across five ancestral populations. We assessed the variability of the total CN by the percentage of individuals having the mode CN. For this study, we say that a gene family has low CN variability if more than 90% of the individuals have the mode CN value. Conversely, a gene family is defined as having high CN variability if less than 90% of individuals have the mode CN value. Based on these definitions, 79 of the gene families have low CN variability and 81 have high CN variability (Figure 2A, Table S1). Additionally, 25.6% (41/160) of the gene families had significant (Chi-squared test, p<0.05, with Bonferroni correction) deviations between ancestral populations (Figure S1).

**Figure 2.**
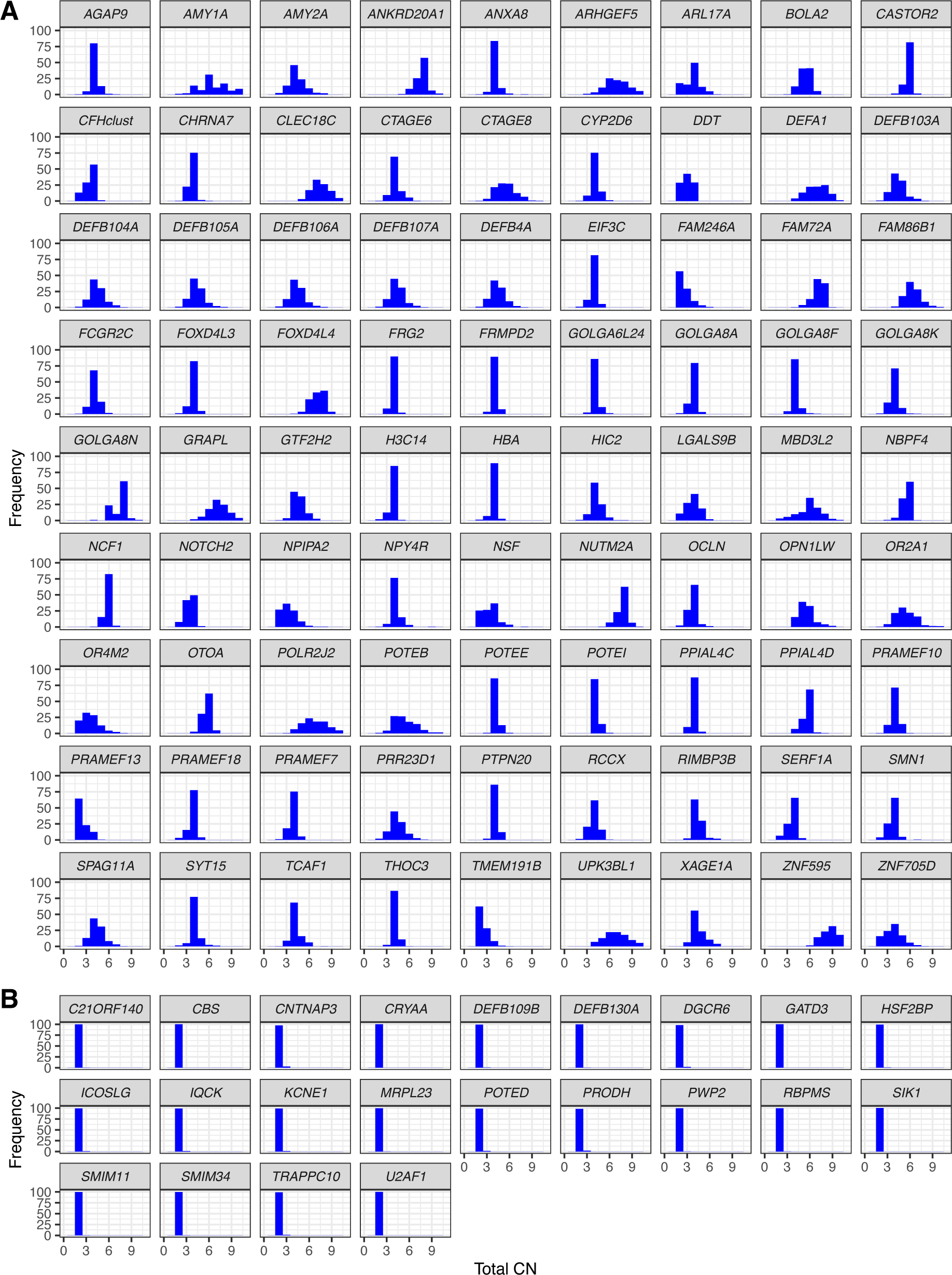
Distribution of the total CN of each gene family across populations. One archetype gene is selected to represent the name of each family. **A**. Gene families with high CN variability. For the two gene families (*OPN1LW* and *XAGE1A*) located on the X chromosome, only female samples are plotted. **B**. False duplication regions in GRCh38, where the majority of individuals have a total CN of two.

The CN variability can give us a general understanding of the population-level “accuracy” of the reference genome (in this case GRCh38). For example, an SD with two homologous regions like *SMN1*/*SMN2* would always have a CN of four in our analysis if the reference is correct and generalizes across the population. Likewise, a gene family where every individual has a CN of two in the population is likely a false SD in the reference. We identified 22 gene families where more than 95% of all individuals have a total CN of two (Table S3, Figure 2B). This suggests that duplications are rare in the population for these genes and these SDs could represent errors in the reference genome. Nineteen of these gene families overlap regions that were classified as false duplications in GRCh38 based on the CHM13 T2T assembly (Nurk et al. 2022). Conversely, three of these gene families (*DEFB109B, CNTNAP3/CNTNAP3C* and *POTED*; see Table S3) were not identified as false duplications in GRCh38 by the CHM13 T2T assembly. However, these three gene families are only present once in the CHM13 T2T assembly. Paraphase analysis of CHM13 data shows one haplotype in *DEFB109B* and *CNTNAP3/CNTNAP3C* but two haplotypes in *POTED*, where the additional haplotype could be due to mosaicism, as the coverage of this region is similar to the genome average (Figure S2).

Additionally, CHM13 has only one gene copy for the *CTAGE8/CTAGE9* family and the *OR2A1/OR2A42* family (Figure S2), resulting in missing genes (*CTAGE8* and *OR2A42*), which were attributed to false duplications in GRCh38 (Nurk et al. 2022). For another gene family, *RIMBP3/RIMBP3B/RIMBP3C*, CHM13 has two copies (Figure S2) and attributed the missing *RIMBP3* to a false duplication in GRCh38. All three of these gene families are truly CN variable regions in the population (distributions shown in Figure 2A) and thus not false duplications in GRCh38.

### Gene families with exceptionally low within-family diversity

Paraphase identified 159,795 haplotypes from the 160 gene families in the 259 samples. Extensive gene conversion and unequal crossing over can result in highly similar gene copies that can no longer be separated into different genes based on sequence alone. For example, *SMN1* and *SMN2* are different in sequence in Exons 7-8, but are indistinguishable in Exons 1-6 indicating that gene conversion is much more common in Exons 1-6 than in Exons 7-8 (Chen et al. 2023). Thus, a principal component analysis (PCA) of variants in Exons 7-8 can differentiate *SMN1* haplotypes from *SMN2* haplotypes, but a PCA of variants in Exons 1-6 does not differentiate the genes (Figure S3).

To identify gene families with low within-family diversity, we developed a metric based on the heterozygosity (divergence) between individual haplotypes (see Methods). For example, in a gene family with a gene and a paralog, the gene will evolve independently from the paralog in the absence of gene conversion. This means that the heterozygosity will be lower between two copies of the gene (i.e. gene-gene heterozygosity) or two copies of the paralog in the absence of any selective pressures. Conversely, the gene-paralog heterozygosity will be significantly higher (Figure S4). Increasing rates of gene conversion and unequal crossing over will tend to make the gene more similar to the paralog and thus drive the gene-paralog heterozygosity down (Figure S4).

We identified 23 gene families (termed low-diversity gene families) where the within-family sequence divergence is comparable to the general allelic sequence divergence (See Methods). Among these, 4 are on chrY, 11 on chrX, and 8 on autosomes (Table 2). It is often not easy to assign haplotypes of a gene family to individual genes without prior knowledge of how genes and paralogs differ from each other. However, among the 23 low-diversity gene families, there are five where the phased haplotypes extend into non-homologous regions so that we can assign haplotypes to genes based on their flanking sequence: *AMY1A/AMY1B/AMY1C* (Figure 3A), *CTAG1A/CTAG1B*, *BOLA2/BOLA2B, SULT1A3/SULT1A4* and *SLX1A/SLX1B* (*BOLA2/BOLA2B, SULT1A3/SULT1A4, and SLX1A/SLX1B* are a group of three families in tandem and genotyped as one region by Paraphase). PCA of the haplotype sequences shows that haplotypes of the different genes of the same family are indistinguishable from each other (Figure 3B-D, also see Figure S5).

**Figure 3.**
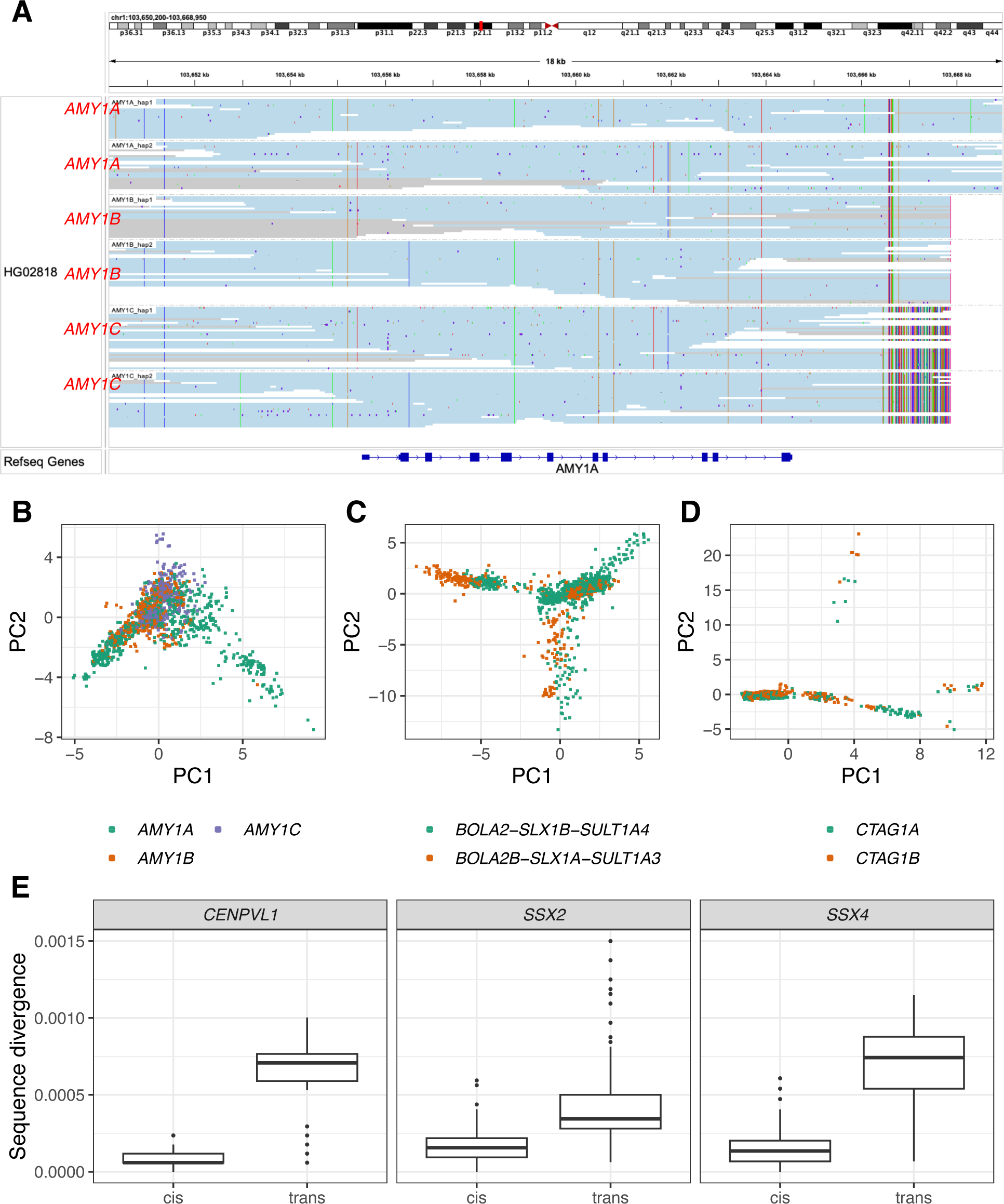
Gene families with low within-family diversity. **A.** Haplotypes of the *AMY1* family in a sample, showing two copies each of *AMY1A*, *AMY1B* and *AMY1C.* Reads in blue are consistent with a single haplotype. Reads in gray are consistent with more than one possible haplotype, i.e. when two or more haplotypes are identical over a region. The ends of the haplotypes extend into downstream non-homologous regions so we can assign the haplotypes into the three genes. **B-D.** PCA of haplotype sequences of the *AMY1A/AMY1B/AMY1C* family (B), *BOLA2-SLX1B-SULT1A4/BOLA2B-SLX1A-SULT1A3* family (a group of three families in tandem and genotyped as one region by Paraphase) (C) and *CTAG1A/CTAG1B* family (D). Each dot represents a haplotype in the population. Colors represent different genes in a family as assigned according to the ending sequences of each haplotype (which extends into non-homologous regions). **E.** Sequence divergence between haplotypes in cis vs. trans in three palindromic gene families. One gene is selected to represent the name of each family (*CENPVL1*: *CENPVL1*/*CENPVL2*. *SSX2*: *SSX2*/*SSX2B*. *SSX4*: *SSX4*/*SSX4B*).

**Table 2.** Gene families with low within-family diversity.

|  | Gene families | Palindromic | High CN variability | Human-specific duplication |
| --- | --- | --- | --- | --- |
| chrY | *BPY2/BPY2B/BPY2C* | x |  |  |
|  | *CDY1/CDY1B* | x |  |  |
|  | *CDY2A/CDY2B* | x |  |  |
|  | *HSFY1/HSFY2* | x |  |  |
| chrX | *CENPVL1/CENPVL2* | x |  |  |
|  | *CTAG1A/CTAG1B* | x |  |  |
|  | *CXorf49/CXorf49B* | x |  |  |
|  | *DMRTC1/DMRTC1B* | x |  |  |
|  | *FAM156A/FAM156B* | x |  |  |
|  | *MAGED4/MAGED4B* | x |  |  |
|  | *NXF2/NXF2B* | x |  |  |
|  | *SSX2/SSX2B* | x |  |  |
|  | *SSX4/SSX4B* | x |  |  |
|  | *TCP11X1/TCP11X2* | x |  |  |
|  | *XAGE1A/XAGE1B* | x | x |  |
| Autosomes | *SLX1A/SLX1B* |  | x | x |
|  | *BOLA2/BOLA2B* |  | x | x |
|  | *SULT1A3/SULT1A4* |  | x | x |
|  | *NPIPA2/NPIPA3* |  | x | x |
|  | *AMY1A/AMY1B/AMY1C* |  | x | x |
|  | *SERF1A/SERF1B* |  | x | x |
|  | *EIF3C/EIF3CL* |  | x |  |
|  | *TRIM49D1/TRIM49D2* | x |  |  |

The 23 low-diversity gene families show two different patterns in their genomic structure, CN variability and evolutionary history (Table 2). Those on autosomes have high CN variability and many are human-specific duplications (See Discussion). Conversely, low-diversity gene families on sex chromosomes mostly have low CN variability, are arranged in palindrome structures and evolutionarily conserved, i.e. all gene family members are present in other primates where they are also in palindromes (Trombetta and Cruciani 2017; Jackson et al. 2021). Additionally, there are 3 palindromic families on chrX where the genes and paralogs are in tandem so we can identify copies on the same chromosome. In these 3 gene families, the gene copies in cis are more similar to each other than those in trans (Figure 3E), suggesting that gene conversion between arms of palindromes happens more frequently in cis (possibly through forming a hairpin structure) than in trans.

### Identification of *de novo* mutations and gene conversion

In 36 parent-offspring trios we identified 12 events (6 paternal and 6 maternal) where a haplotype in the proband is different from the corresponding haplotype in the parent (Figure S6-S16). Eleven of these are *de novo* events where the proband haplotype differs from the parent haplotype by one SNV. Among these, 7 are *de novo* SNVs and 4 are products of gene conversion. Among the gene conversion cases, 2 are non-allelic (an example is shown in Figure 4), 1 is allelic and 1 could be either allelic or non-allelic. Among the 11 *de novo* events, 4 are intergenic, 6 are in introns, and 1 is in an exon (synonymous). The remaining case of the 12 events is a hybrid haplotype between two haplotypes from the same parent, which could arise through equal or unequal crossing over (inconclusive without longer range phasing information in the parent due to the high copy number of the gene family) (Figure S16).

**Figure 4.**
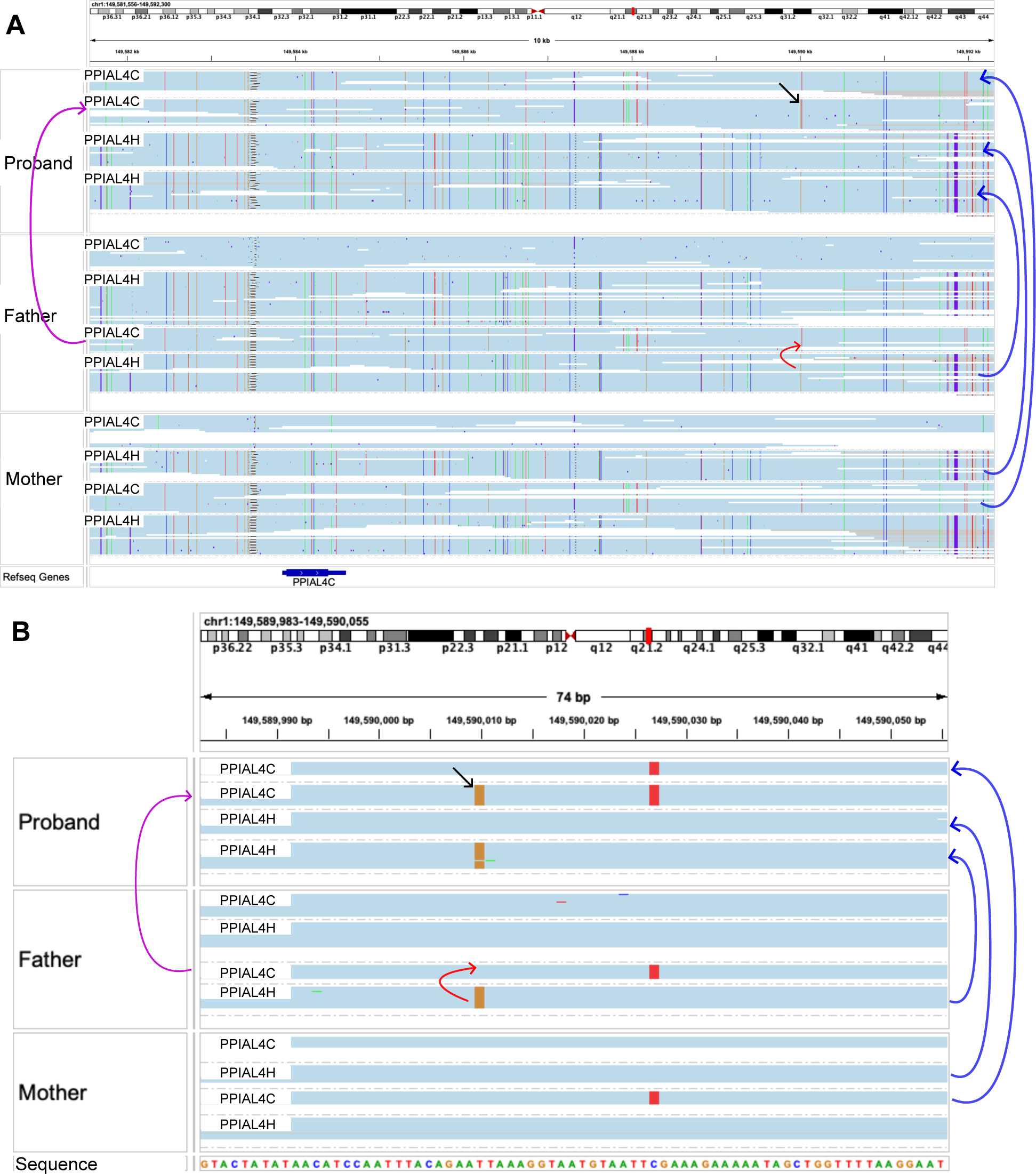
*De novo* non-allelic gene conversion in a trio. **A.** The black arrow denotes the SNV created by non-allelic gene conversion on haplotype 2 of the proband. It is not present on the inherited haplotype, haplotype 3 in the father. Instead, it is present on haplotype 4 of the father, which belongs to the other gene in the family. The magenta arrow on the left indicates the inheritance of the converted haplotype. The blue arrows on the right indicate the inheritance of the other haplotypes. The red arrow shows the direction of the gene conversion. **B.** Close view of the converted variant.

### Resolving medically relevant homologous genes

As a demonstration of how Paraphase can be used to study gene families in the population, we examined variant and haplotype frequencies across populations in three known medically relevant gene families, *CYP21A2/CYP21A1P*, *PMS2/PMS2CL* and *OPN1LW*/*OPN1MW*.

Variants in *CYP21A2* cause 21-Hydroxylase-Deficient Congenital Adrenal Hyperplasia (21-OHD CAH) (Merke and Auchus 2020). *CYP21A2* resides in a 30-kb tandem repeat called the RCCX module that includes its pseudogene, *CYP21A1P*, together with two other pairs of paralogs, *C4A/C4B* and *TNXB/TNXA* (Pignatelli et al. 2019; Merke and Auchus 2020) (Figure 5A). This region is susceptible to gene conversion (Merke and Auchus 2020), as well as deletions and duplications of the RCCX module resulting in CN changes and disease-causing hybrid genes between *CYP21A2* and *CYP21A1P*. The total CN of RCCX is highly variable across populations (Figure 5B) with 38.2% of individuals having a CNV. Figure 5A shows examples of samples with various CNs. In addition, we identified a duplication allele (Figure 5A, bottom panel) that carries a copy of *CYP21A1P*, a copy of *CYP21A2* with a stop-gain variant Q319X, and a second functional copy of *CYP21A2*. This allele is found at 1-2% frequency in the populations (Table S4) and, without phasing the full region, could be misidentified as a pathogenic allele due to the presence of Q319X. Researchers have previously found that individuals with Q319X frequently have a duplication of *CYP21A2*, which complicates *CYP21A2* testing (Lekarev et al. 2013). Paraphase can distinguish a *CYP21A2*+*CYP21A2*(Q319X) allele vs. a *CYP21A2*(Q319X) allele.

**Figure 5.**
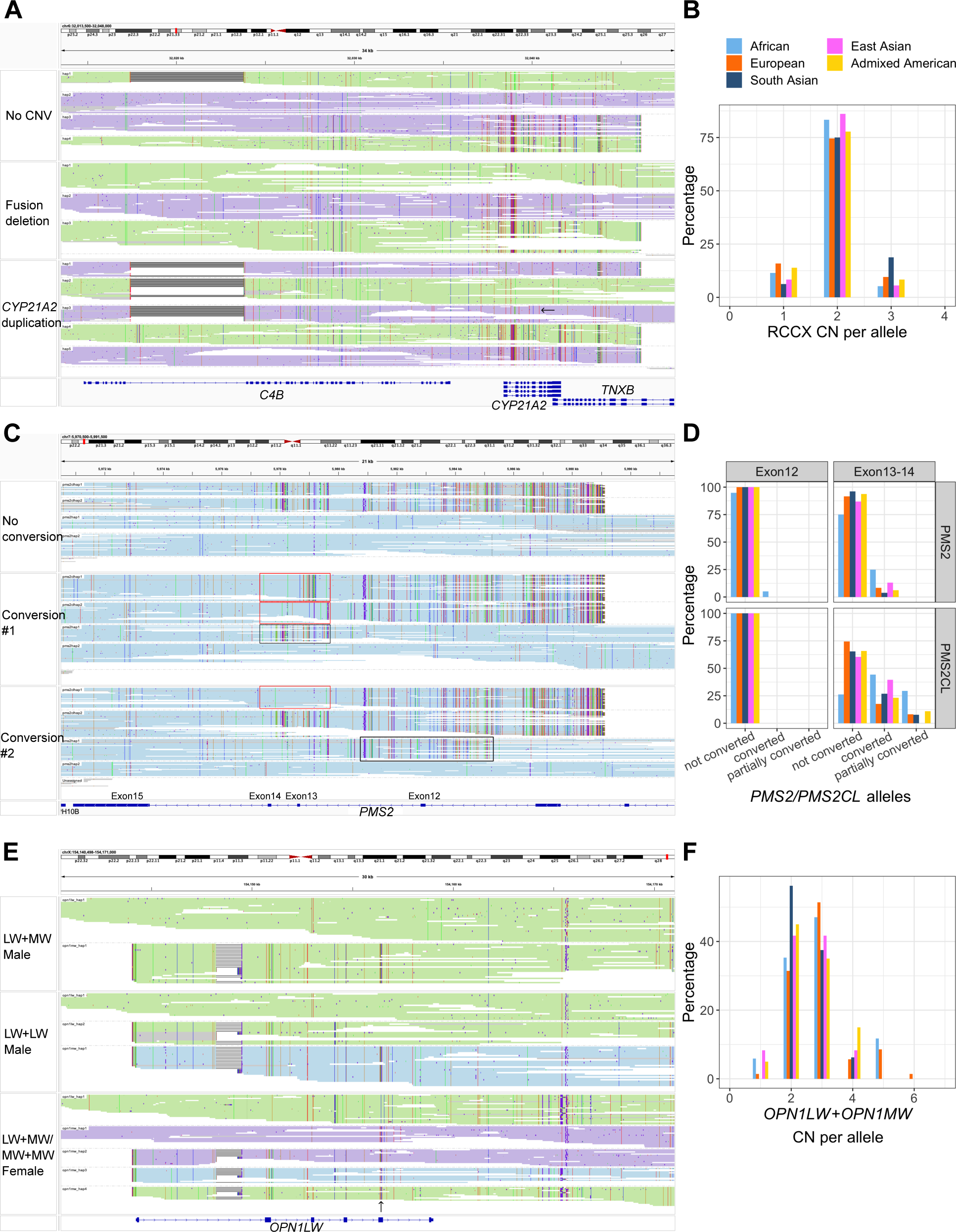
Population results in *CYP21A2*, *PMS2* and *OPN1LW/OPN1MW*. **A.** Paraphase resolved haplotypes in the RCCX module, realigned to the RCCX copy that encodes *CYP21A2*. Haplotypes of the same color (purple or green) are from the same allele. Longer haplotypes represent the last RCCX copies in the array on each allele, and shorter haplotypes represent remaining copies. Three examples are shown, including a sample with no CNV (top), a sample with fusion deletion (*CYP21A1P*-*CYP21A2* fusion, middle, purple allele) and a sample with RCCX duplication (bottom). The bottom sample carries an allele (purple) with a wild-type copy of *CYP21A2* and another copy of *CYP21A2* harboring a pathogenic variant Q319X (black arrow). **B.** Frequency of the total RCCX CN per allele across populations. **C.** Paraphase resolved haplotypes in *PMS2*/*PMS2CL*, realigned to *PMS2*. Exon numbers are labeled with respect to *PMS2*. Longer haplotypes represent *PMS2* copies and shorter ones represent *PMS2CL* copies. Three examples are shown, including a sample with no gene conversion and two samples carrying alleles converted in Exon12 or Exon13-14 (conversions in *PMS2* shown in black boxes and conversions in *PMS2CL* shown in red boxes). **D.** Frequency of gene conversions between *PMS2* and *PMS2CL* across populations in Exon12 and Exon13-14. **E.** Paraphase resolved haplotypes in *OPN1LW*/*OPN1MW*, realigned to *OPN1LW*. Haplotypes of the same color (purple or green) are from the same allele. Longer haplotypes represent the first copies of the repeat in the array on each allele, and shorter haplotypes represent remaining copies. The blue color indicates gene copies beyond the second copy in the array, i.e. not expressed. *OPN1LW* and *OPN1MW* are assigned based on variants in Exon5 (black arrow). Three examples are shown, including a normal allele with a copy of *OPN1LW* followed by a copy of *OPN1MW* (top), an allele (*deutan*) with a copy of *OPN1LW* followed by a copy of *OPN1LW* (middle, the third unexpressed copy marked in blue), and a female sample where one normal allele has a copy of *OPN1LW* followed by a copy of *OPN1MW*, and the other allele (*protan*) has a copy of *OPN1MW* followed by a copy of *OPN1MW* (bottom, the third unexpressed copy marked in blue). **F.** Distribution of the summed CN of *OPN1LW* and *OPN1MW* per allele across populations.

Pathogenic variants in *PMS2* cause Lynch syndrome (ten Broeke et al. 2018). In its last few exons (Exons 12-15), *PMS2* has high sequence similarity to its pseudogene *PMS2CL*, and gene conversion and unequal crossing overs are known to promote sequence exchange between the two genes (Hayward et al. 2007; van der Klift et al. 2010; Ganster et al. 2010). In Exon 15, the sequences of *PMS2* and *PMS2CL* are indistinguishable from each other, lacking any differentiating variants (Figure 5C, also see PCA in Figure S17). For example, a commonly considered *PMS2CL*-specific variant (Wernstedt et al. 2012), NM_000535.7:c.*92dup, is in 72.8% of *PMS2CL* alleles and 31.1% of *PMS2* alleles. We found that gene conversion happens between *PMS2* and *PMS2CL* occasionally in Exon 12 and frequently in Exons 13-14 (See Methods and Figure 5C-D). Interestingly, our analysis shows more evidence of gene conversion in individuals of African ancestry, and more than 75% of African *PMS2*/*PMS2CL* alleles are partially or fully converted (Figure 5D).

*OPN1LW* and its paralog *OPN1MW* are responsible for red-green color vision deficiencies and other vision conditions such as blue cone monochromacy (BCM) (Neitz and Neitz 2011). The region is arranged in a gene array and only the first two genes in the array are expressed (Neitz and Neitz 2011). Paraphase identifies all copies of the repeat, assigns genes to *OPN1LW* or *OPN1MW*, and identifies the first two copies in the array on each chromosome. Figure 5E shows alleles with one copy each of *OPN1LW* and *OPN1MW* (top panel), as well as alleles that only have *OPN1LW* (middle panel) or *OPN1MW* (bottom panel) in the first two copies of the array. The total CN of this array is highly variable among populations (Figure 5F). Allele frequencies are summarized in Table S5, including alleles that cause color vision deficiencies.

In addition to the three gene families described above, we also summarized population results for other medically relevant genes, including *SMN1/SMN2* (spinal muscular atrophy (Lunn and Wang 2008), Figure S18), *STRC* (hereditary hearing loss and deafness (Yokota et al. 2019), Figure S19), *HBA1/HBA2* (Alpha thalassemia (Farashi and Harteveld 2018), Table S6), *IKBKG* (Incontinentia Pigmenti (Cammarata-Scalisi et al. 2019), Table S7), the *CFH* gene cluster (*CFH/CFHR1/CFHR2/CFHR3/CFHR4*) (atypical hemolytic uremic syndrome (Zipfel et al. 2007) and age-related macular degeneration (Klein et al. 2005; Hughes et al. 2006), Table S8) and *GBA* (Gaucher and Parkinson’s disease (Hruska et al. 2008; Sidransky et al. 2009), Table S9). Together, we identified medically relevant variants in these 9 gene families in 75%, 49.6%, 45.8%, 52.2% and 17.4% of individuals of African, European, Admixed American, South Asian and East Asian ancestries, respectively.

## Discussion

In this paper, we applied Paraphase to 160 segmental duplication regions where large (>10kb) regions of high (>99%) sequence similarity exist between genes and their paralogs. By phasing reads from the same gene family together, Paraphase can recover misaligned reads and correctly resolve genes together with their highly similar paralogs/pseudogenes. This method enables high-throughput CN detection and genotyping of SD-encoded genes with only HiFi data at standard WGS depth (30X).

An important benefit of the gene-family-centered approach is that it is not influenced by the CN difference between an individual and the reference, and it can work even when the CN of a gene family does not agree with the reference genome in most individuals of the population, such as in the case of false segmental duplications in GRCh38. In addition, by calling variants against the same reference gene within a gene family, Paraphase outputs gene copies that can be easily compared against each other, allowing us to perform within-family divergence analysis, as well as to detect de novo mutations including gene conversion events between paralogs. Our analysis of 36 trios identified 7 *de novo* SNVs and 4 *de novo* gene conversion events, demonstrating the power of long-read sequencing in detecting de novo variations (Noyes et al. 2022; Kucuk et al. 2023), particularly in previously inaccessible regions of the genome.

Among regions analyzed by Paraphase, we observed that gene families on sex chromosomes are more CN invariable (93.1% of gene families on sex chromosomes have low CN variability vs. 39.7% on autosomes) and have drastically lower within-family diversity (median pairwise haplotype divergence 0.00033 on sex chromosomes vs. 0.00187 on autosomes, p-value 4.179e-11). This could be related to the fact that most gene families on sex chromosomes are arranged in palindrome structures (86.2% vs. 16.8% on autosomes). Unequal crossing overs between arms of a palindrome result in inversions and do not change the copy number. Arm-to-arm gene conversion is known to occur frequently to prevent sex chromosomes from accumulating deleterious mutations in the absence of homologous chromosomes (Swanepoel et al. 2020; Jackson et al. 2021), and could contribute to the low within-family diversity.

We identified 23 gene families with extremely low within-family diversity (Table 2), where extensive gene conversion and/or unequal crossing over have resulted in paralogs that are as similar as alleles from the same gene. Consistent with most other families on sex chromosomes, the low-diversity gene families on sex chromosomes are all arranged in palindromes and mostly have low CN variability. For these gene families, both the genes and the palindrome structure are evolutionarily conserved in other primates. The low-diversity gene families on autosomes, however, are not arranged in palindrome structure and mostly have high CN variability. Interestingly, many of these gene families are duplicated exclusively in the human lineage, with positive selection detected, e.g. *AMY1A*/*AMY1B*/*AMY1C* (Perry et al. 2007), *BOLA2/BOLA2B* (Nuttle et al. 2016; Giannuzzi et al. 2019) and *SULT1A3/SULT1A4* (Butcher et al. 2018). It is possible that extensive gene conversion and unequal crossing over happens between members of these families to prevent the newly duplicated genes from diverging to maintain an elevated gene dosage in humans. For example, the CN of the salivary amylase gene (*AMY1A*/*AMY1B*/*AMY1C*, Figure 3A-B) is variable between populations (Figure S1) and is correlated with salivary amylase level (Perry et al. 2007). *AMY1* is under positive selection so that populations with high-starch diets have more copies of *AMY1*, which improves the digestion of starchy foods (Perry et al. 2007). Beyond gene families with low within-family diversity throughout the entire gene body, one future direction is to extend this analysis to identify local low-diversity regions resulting from gene conversions, such as the gene conversion found in Exons 1-6 of *SMN1/SMN2* (Figure S3), and Exon 15 of *PMS2* (Figure 5C, Figure S17).

The SD-encoded genes presented in this paper were previously inaccessible to population-wide genomic analyses and hence are largely missing from variant annotation databases such as gnomAD (Karczewski et al. 2020), creating hurdles in variant interpretation. Here we provide a database (https://zenodo.org/doi/10.5281/zenodo.10909886) of variant allele frequencies collected from the population samples used in this paper. This annotation resource can be further expanded as more HiFi data are generated and analyzed with Paraphase.

One limitation of Paraphase is that currently it focuses on gene families with <10 copies in an individual and does not include other highly similar genes with even higher CNs. This excludes 79 genes that fall into SDs in our analysis. Nevertheless, Paraphase can be customized to analyze user-specific regions, allowing new targets to be added in the future.

Paraphase, combined with HiFi long reads, provides a single framework for resolving paralogous genes. In medically important genes challenged by pseudogenes or paralogs, Paraphase helps enable more accurate testing to detect pathogenic variants, thus bringing us one step closer to consolidating the numerous currently offered genetic tests into a single test. Furthermore, in previously inaccessible and less studied genes, population-wide sequencing based analysis with Paraphase will facilitate the discovery of novel gene-disease associations.

## Methods

### Paraphase: HiFi-based caller for highly similar paralogous genes

Paraphase is designed to work with both PacBio HiFi WGS and target enrichment data. Paraphase resolves a group of highly similar genes by extracting HiFi reads aligned to any member of the family, realigning them to the archetype gene, and phasing them into haplotypes, followed by variant calling on each haplotype (Chen et al. 2023) (Figure 1A). Realigning all reads from all genes of the same gene family to one gene bypasses the error-prone process of aligning reads to multiple similar regions. This framework enables all copies of the gene family, including genes and their paralogs or pseudogenes, to be examined for variants and annotated for functional status.

When two homologous regions are in tandem, Paraphase uses read-based phasing to further phase gene haplotypes into alleles, i.e. gene copies on the same chromosome, by grouping haplotypes that have an overlapping set of supporting reads. For example, for the RCCX module demonstrated in Figure 5A, reads are grouped by the haplotypes they originate from and haplotypes of the same color (green or purple) represent those from the same allele.

Gene fusions between paralogs are called by detecting haplotypes whose flanking regions (upstream and downstream of the homologous region) are consistent with two different genes. Fusion breakpoints are called by detecting a switch in bases at the paralogous sequence variant (PSV) sites.

Within Paraphase, there are a few gene-specific callers for medically relevant genes. These callers use gene-specific information during analysis, for example, known sequence differences between genes and paralogs/pseudogenes. In addition, these callers produce gene-specific output information such as hybrid gene structures and known pathogenic variants, including large difficult-to-call structural variants.

### Genome-wide identification of highly similar genes for analysis by Paraphase

We extracted 19,394 Ensembl protein coding genes (>20 kb sequences centered on each gene, adding flanking sequences for shorter genes) and aligned them against GRCh38 (ALT contigs excluded, pseudoautosomal regions (PARs) masked). We selected genes that had alignment matches >10 kb in length and >99% in sequence similarity as candidate gene families. The majority of genes have zero paralogs, and the remaining ones vary in the number of paralogs (Figure S20). Among genes with three or fewer paralogs, which represent the majority of genes with paralogs, we incorporated into Paraphase 155 groups of regions. In addition, we included genes impacted by shorter homology or lower sequence similarity, where gene fusions are highly medically relevant yet difficult to call by conventional SV callers due to homology, including *HBA1/HBA2* (Alpha thalassemia), *GBA1/GBAP1* (Gaucher and Parkinson’s disease), *CYP2D6/CYP2D7* (pharmacogenomics), *CYP11B1/CYP11B2* (Glucocorticoid-remediable aldosteronism) and *CFH/CFHR1/CFHR2/CFHR3/CFHR4* (atypical hemolytic uremic syndrome and age-related macular degeneration). In total, Paraphase analyzes 160 groups of regions (Table S1), which encode 316 genes in total (pseudogenes are not included).

### MAPQ distributions

We selected 20 samples from five ancestral populations with both Illumina (data downloaded from the 1000 Genomes Project (The 1000 Genomes Project Consortium 2015)) and HiFi WGS data available to assess alignment MAPQs. For each gene family of interest, we calculated the median values across the MAPQs of all reads aligned to the homology region in any gene of the gene family.

### Validation

Validation samples were collected from 21 clinical samples (disease or carrier samples) with 30 pathogenic variants in 8 disease-causing genes that were previously validated by orthogonal methods, such as MLPA and Sanger sequencing (Table 1 and Table S2). In addition, we used 36 trios to examine the consistency of haplotypes called in probands vs. parents. Among the 36 trio, 8 were collected from Radboud University Medical Center, 10 were from the 100,000 Genomes Project and 18 were from Genomics Research to Elucidate the Genetics of Rare Diseases (GREGoR) Consortium.

### Population samples

For frequency calculations, we used 259 unrelated individuals from five ancestral populations (113 Europeans, 52 Africans, 48 Admixed Americans, 23 South Asians and 23 East Asians), collected from the Human Pangenome Reference Consortium (HPRC) (Wang et al. 2022; Liao et al. 2023), the 100,000 Genomes Project, Radboud University Medical Center, Genomics Research to Elucidate the Genetics of Rare Diseases (GREGoR) Consortium, and Genomic Answers for Kids (GA4K) at Children’s Mercy Kansas City.

### Gene families with low within-family diversity

We searched for gene families within which the haplotype diversity is comparable to the general sequence diversity between alleles of the same gene. To profile the average allelic sequence divergence, we used Paraphase to phase haplotypes sequences for 600 randomly selected genes (400 on autosomes and 200 on chrX) that fall outside of SDs, i.e. each individual is expected to have two copies of a gene when there is no CNV, in the same set of 259 individuals. Focusing only on SNVs in non-homopolymer regions, 90% of the sequence divergence among haplotypes were lower than 0.00156 for autosomal genes (Figure S21), and for chrX genes, 90% of the sequence divergence among haplotypes were lower than 0.00101 (Figure S21), consistent with a lower mutation rate on chrX (Hodgkinson and Eyre-Walker 2011). To obtain candidate gene families where the within-family sequence divergence is as low as the general allelic diversity, we required that 90% of the sequence divergence among haplotypes of the same gene family to be lower than 0.00156 and 0.00101 for autosomal and sex chromosome gene families, respectively. For both randomly selected genes and Paraphase gene families, only haplotypes from the same ancestral populations were compared for pairwise divergence calculation. Between a pair of haplotypes, sequence divergence was calculated by dividing the number of base differences by the length of the region. For gene families reported in Table 2, we further filtered out gene families where the homology does not span the entire gene, i.e. partial paralogs. Principal component analysis (PCA) within a gene family was conducted on SNV sites identified across all haplotypes of the gene family using the prcomp function in R.

### *PMS2* gene conversion calling

*PMS2* and *PMS2CL* haplotype sequences in Exon 12 region (chr7:5,981,000-5,985,000, GRCh38) and Exons 13-14 region (chr7:5,977,000-5,980,000) are separated into two main groups (*PMS2*-like and *PMS2CL*-like) based on the PCA (Figure S17). Variants (called against the *PMS2* reference sequence) that are present in >95% of the *PMS2CL*-like group and <5% of the *PMS2*-like group are selected as signature sites for calling gene conversion. Gene conversion is called when a *PMS2CL* haplotype has <20% of the signature variants or when a *PMS2* haplotype has >80% of the signature variants. A partial conversion in Exons 13-14 is a special haplotype common in the population, called based on a predefined subset of the signature variants (Figure 5C, middle panel, first haplotype).

## Data access

Paraphase can be downloaded from https://github.com/PacificBiosciences/paraphase HPRC HiFi data can be downloaded from https://github.com/orgs/human-pangenomics/repositories

## Acknowledgements

We thank John Belmont for his valuable comments for the manuscript. We acknowledge colleagues from the diagnostic division of the Human Genetics Department from Radboudumc (Genome Diagnostics Nijmegen) and the Radboud Genomics Technology Center for their technical assistance and the library preparation and sequencing of all samples provided from Radboudumc, in particular Ronny Derks, Amber den Ouden, Janneke Weiss, and Lonneke Haer-Wigman. We thank the Human Pangenome Reference Center (HPRC) for generating and releasing the HiFi WGS data. We thank the Genomic Answers for Kids (GA4K) program at Children’s Mercy Kansas City for generating the GA4K HiFi sequencing data. This research was made possible through access to the 100,000 Genomes Project data in the National Genomic Research Library, which is managed by Genomics England Limited (a wholly owned company of the Department of Health and Social Care). The National Genomic Research Library holds data provided by patients and collected by the NHS as part of their care and data collected as part of their participation in research. The National Genomic Research Library is funded by the National Institute for Health Research and NHS England. The Wellcome Trust, Cancer Research UK and the Medical Research Council have also funded research infrastructure.

## Declaration of interest

X.C., D.B., E.D. and M.A.E. are employees of PacBio. J.M.D., J.N., A.S.B., R.B., K.S.H., L.L., P.K. and S.N. are employees of GeneDx.

## Supplemental information

### Supplementary figures

**Figure S1.**
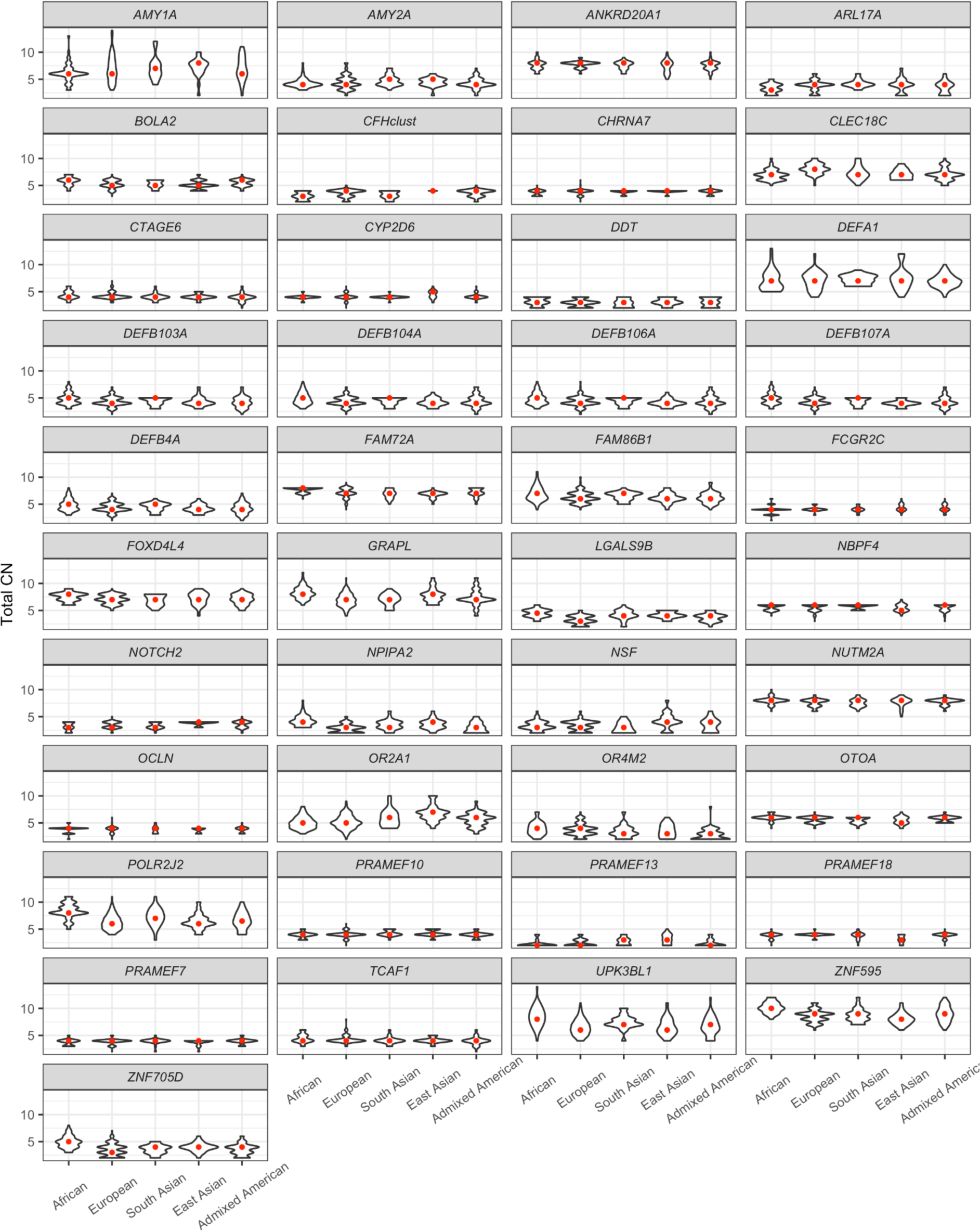
Gene families with significant CN deviations across populations. One archetype gene is selected to represent the name of each family.

**Figure S2.**
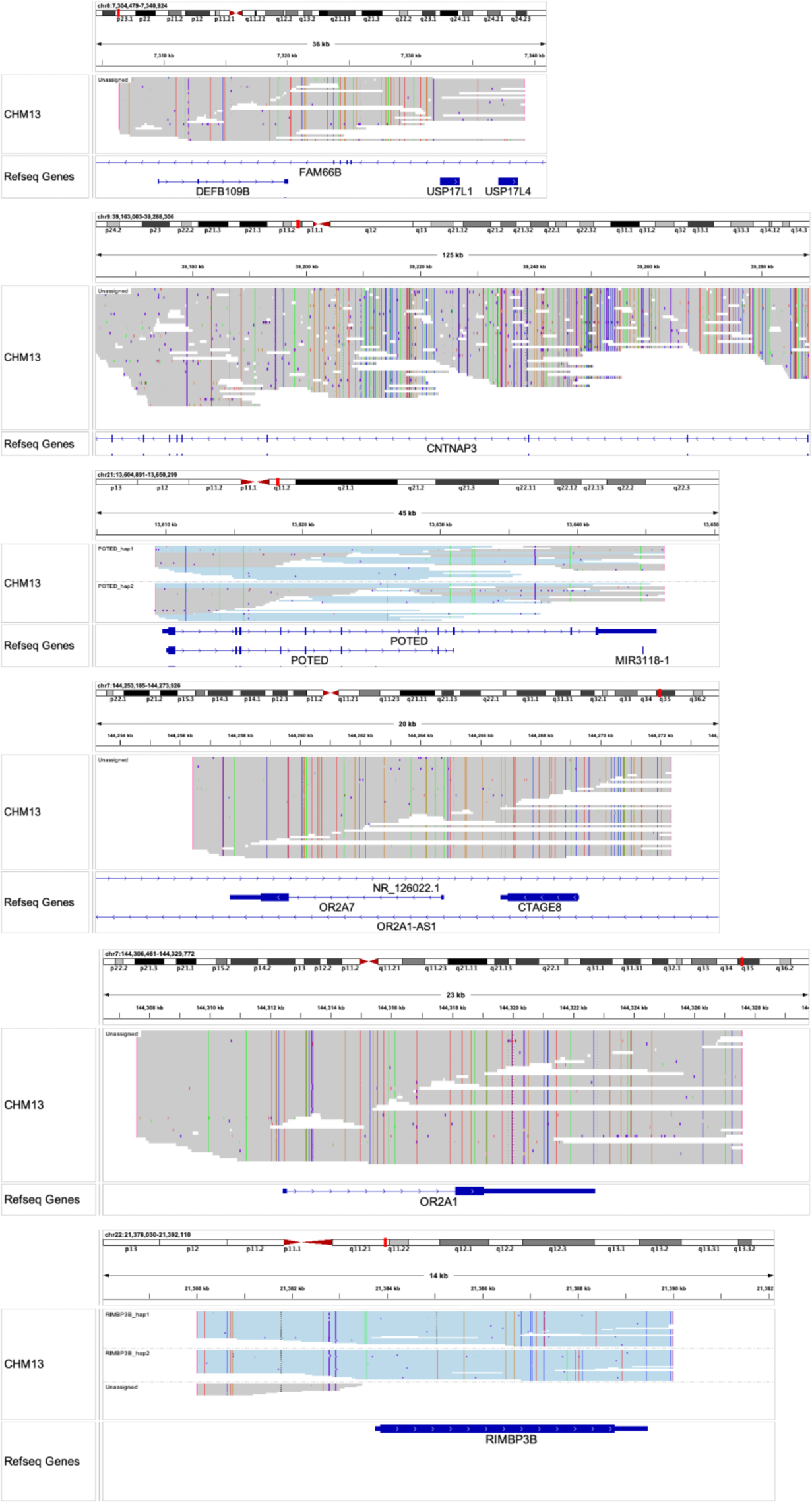
Paraphase results in false duplication regions with CHM13 data. First three: false duplication regions not previously found in CHM13 T2T assembly (*DEFB109B*, *CNTNAP3/CNTNAP3C* and *POTED*). Last three: regions erroneously assigned to false duplications by CHM13 T2T assembly (*CTAGE8/CTAGE9*, *OR2A1/OR2A42* and *RIMBP3/RIMBP3B/RIMBP3C*).

**Figure S3.**
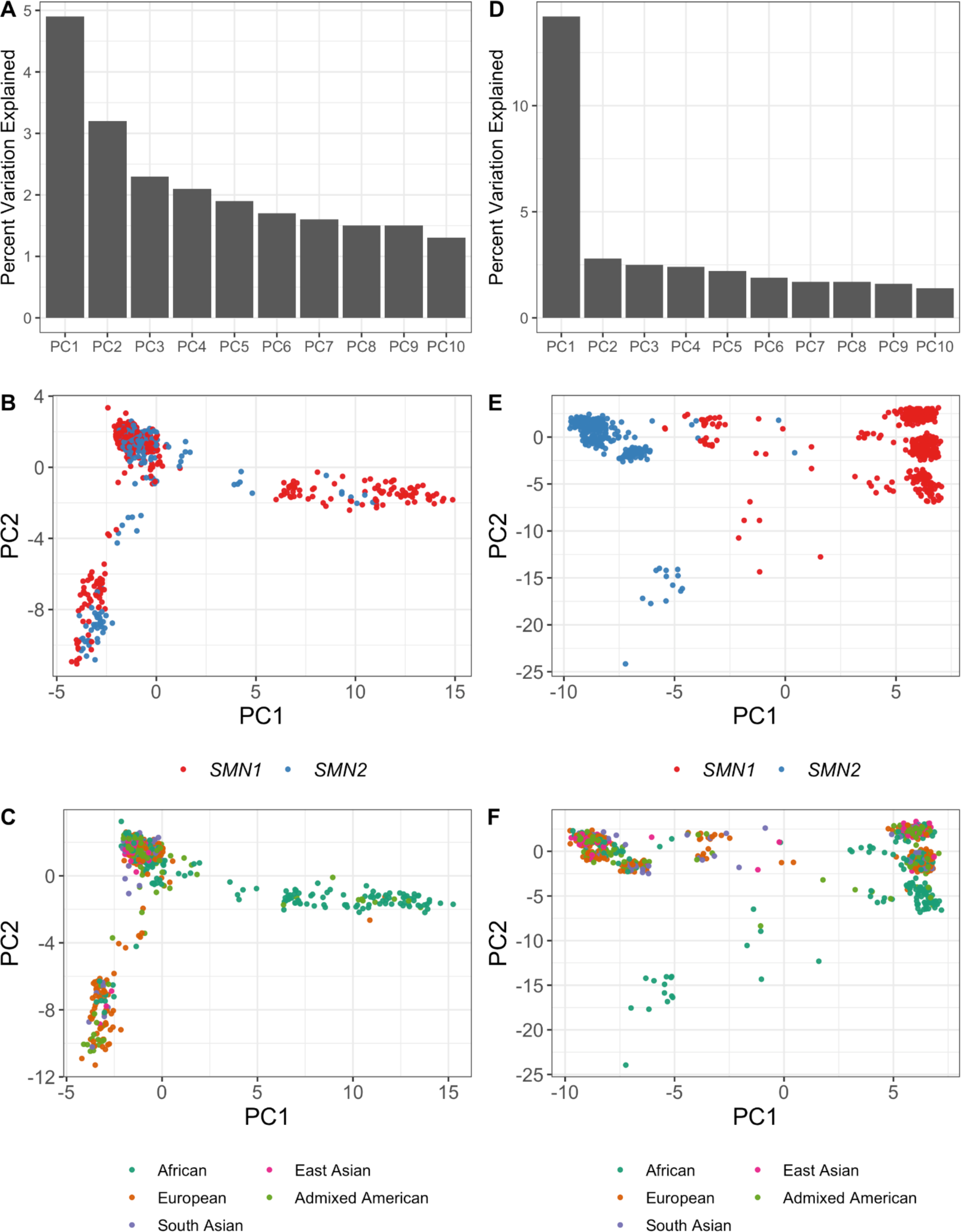
Principal component analysis (PCA) of Exons 1-6 and Exons 7-8 sequences in *SMN1/SMN2*. Exons 1-6 (A-C) sequences are highly similar to each other, with little variation explained by each PC (A). *SMN1* and *SMN2* haplotype sequences do not separate into distinct clusters (B). Instead, the separation on PC1 is by ancestry (C). Exons 7-8 (D-F) are more differentiated, with a much higher level of variation explained by PC1 (D), which separates haplotype sequences by gene (E) rather than ancestry (F).

**Figure S4.**
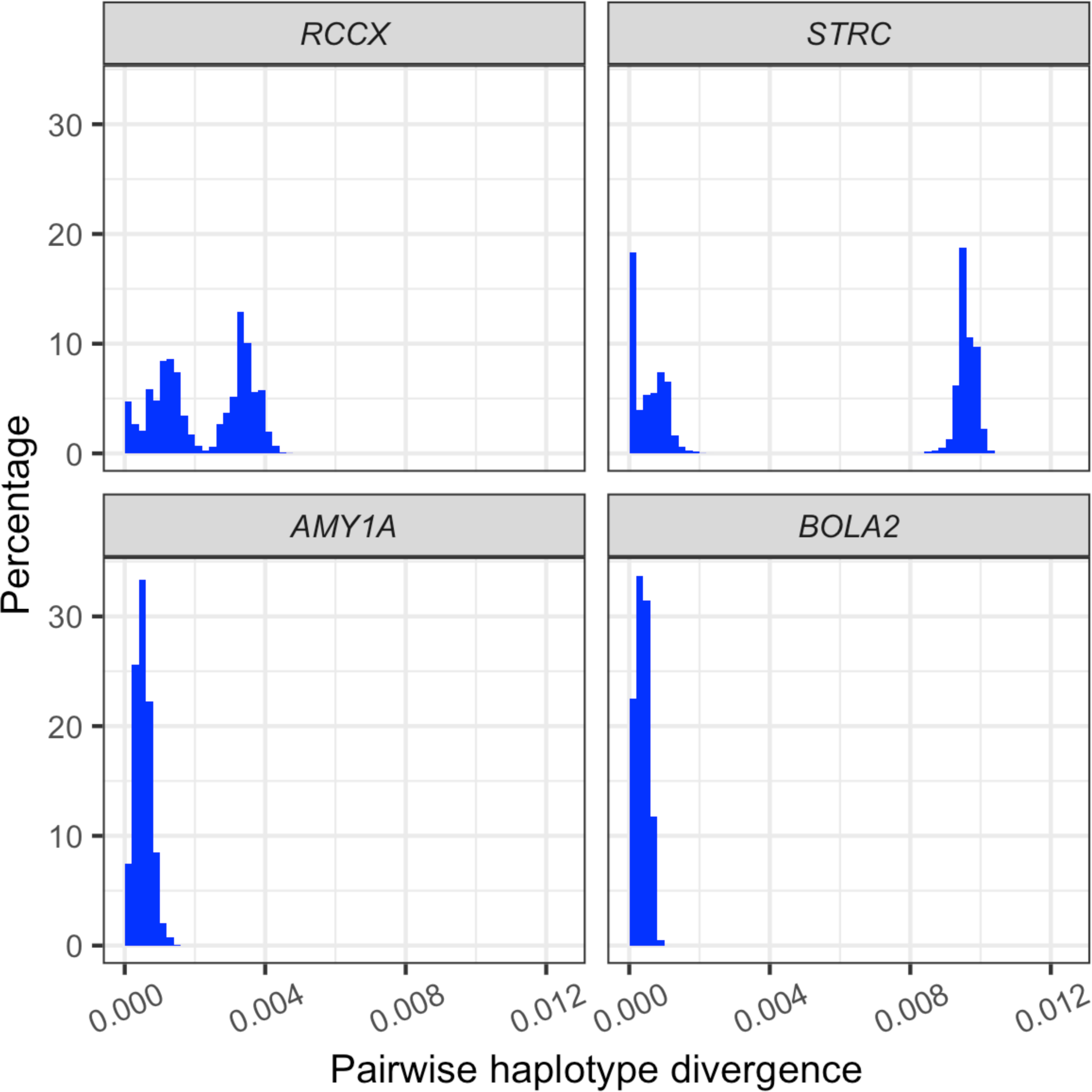
Histograms of pairwise sequence divergence for gene families with differentiated genes and paralogs (upper panel) vs. gene families with low within-family diversity (lower panel). With differentiated genes in the same family, we expect to see two distinct peaks representing low gene-gene (and paralog-paralog) sequence divergence and high gene-paralog sequence divergence. On the other hand, in gene families with low within-family diversity, gene-gene (and paralog-paralog) sequence divergence is similar to gene-paralog sequence divergence and they do not form two peaks. RCCX contains *CYP21A2/CYP21A1P*, *C4A/C4B* and *TNXB/TNXA*.

**Figure S5.**
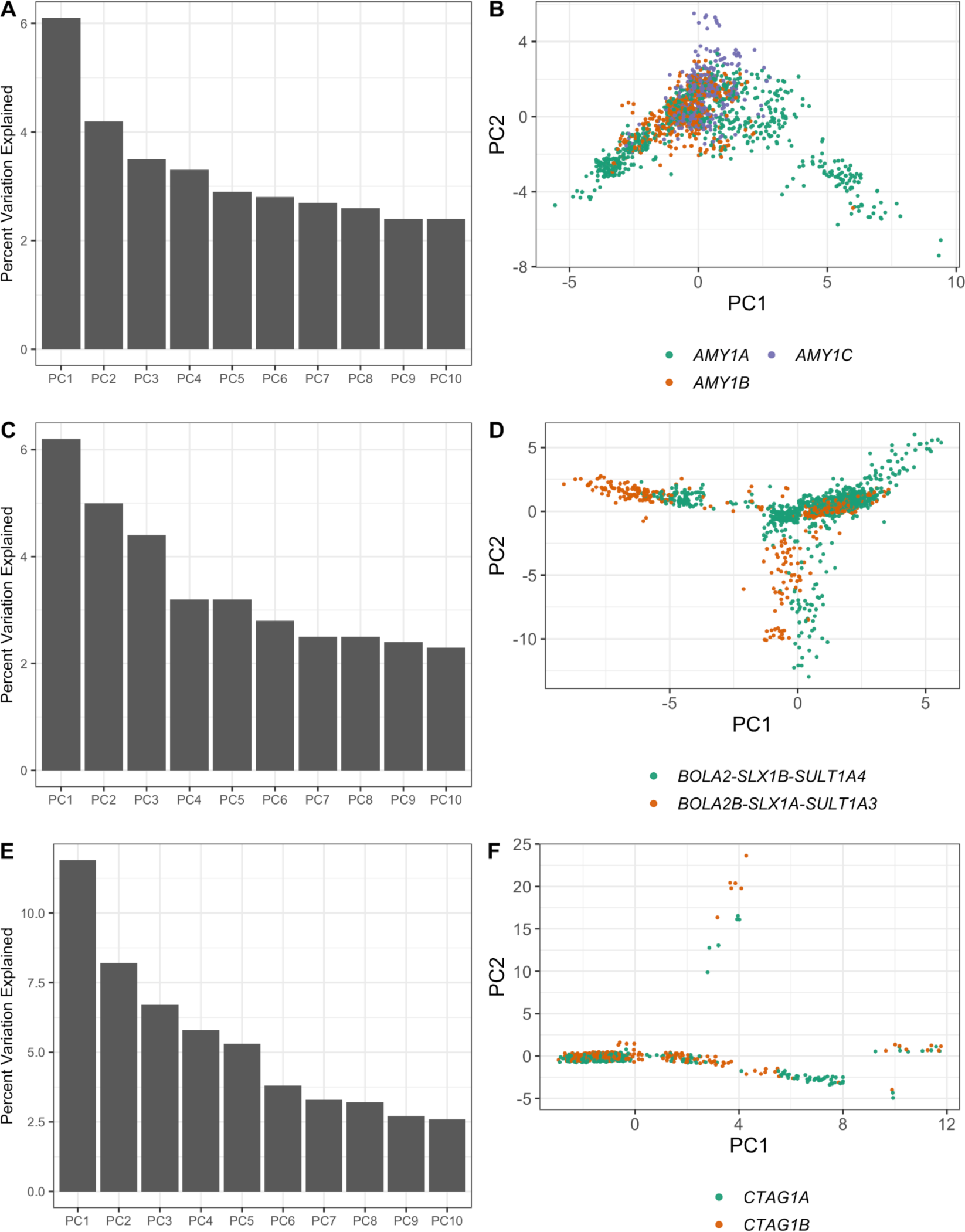
Details of PCA results shown in Figure 3B-D. Each row shows the PCA results for one gene family.

**Figure S6.**
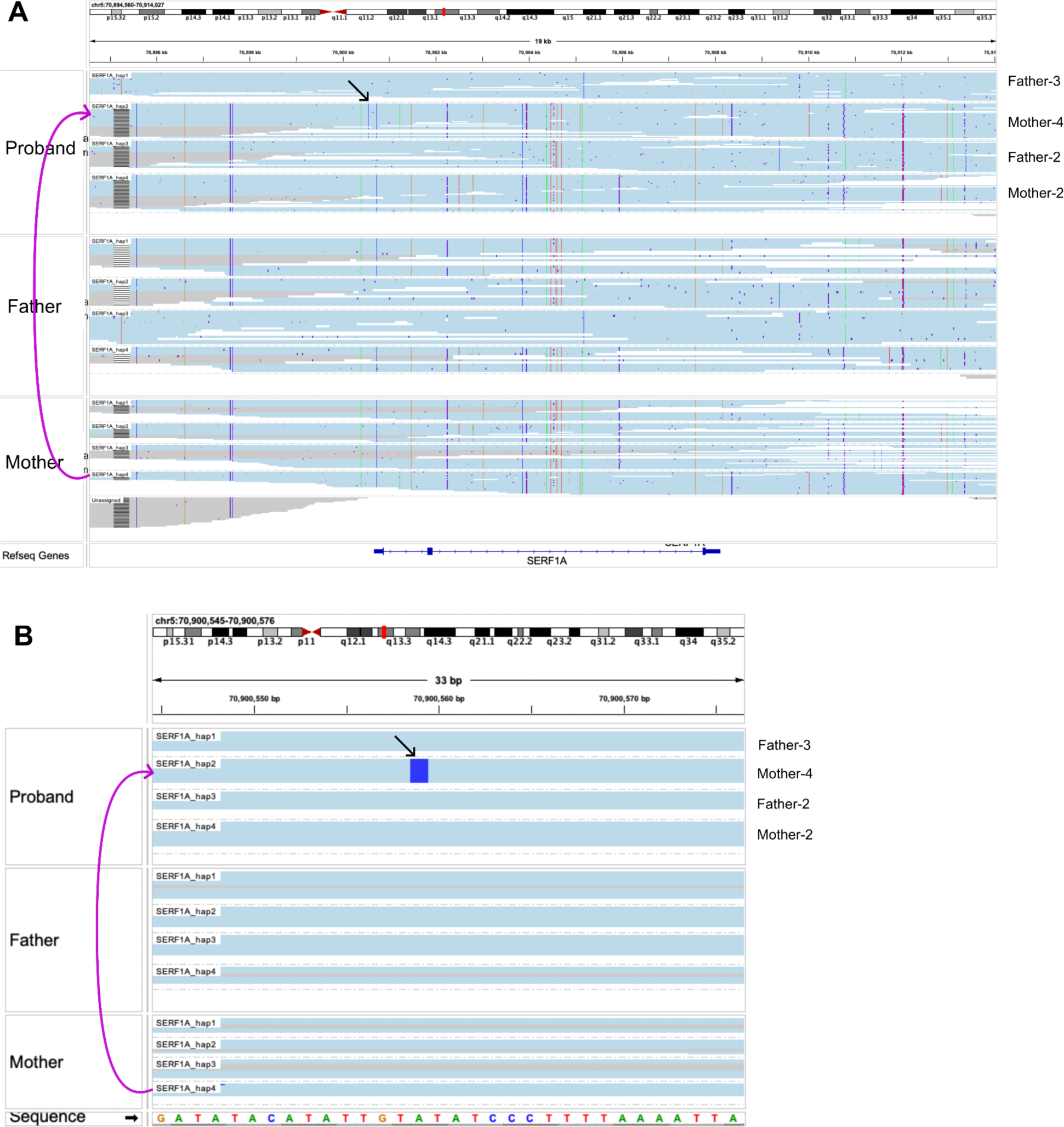
*De novo* SNV #1. Variant is intergenic. Variant is marked by the black arrow. B is a close view of the variant.

**Figure S7.**
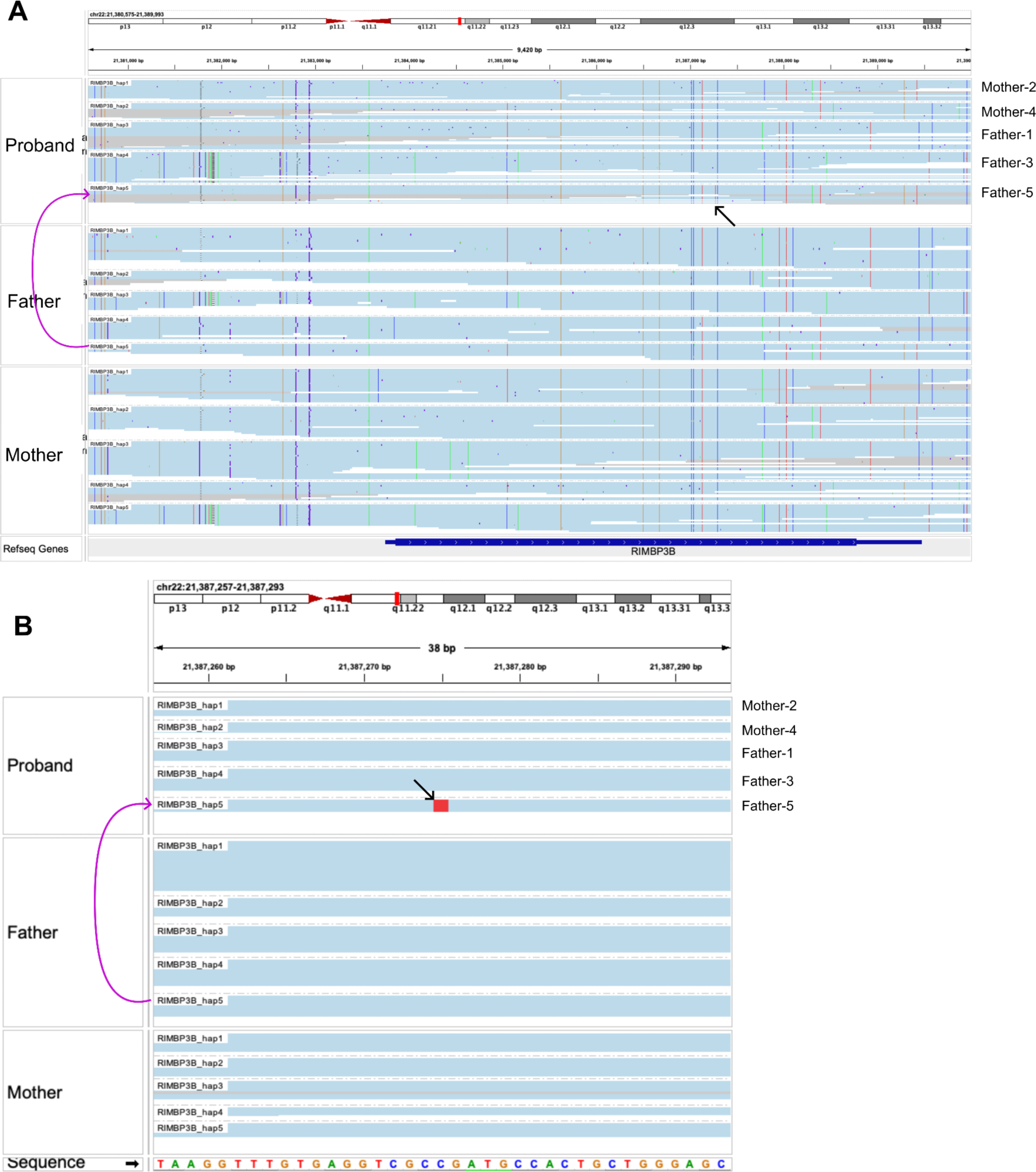
*De novo* SNV #2. Variant is synonymous. Variant is marked by the black arrow. B is a close view of the variant.

**Figure S8.**
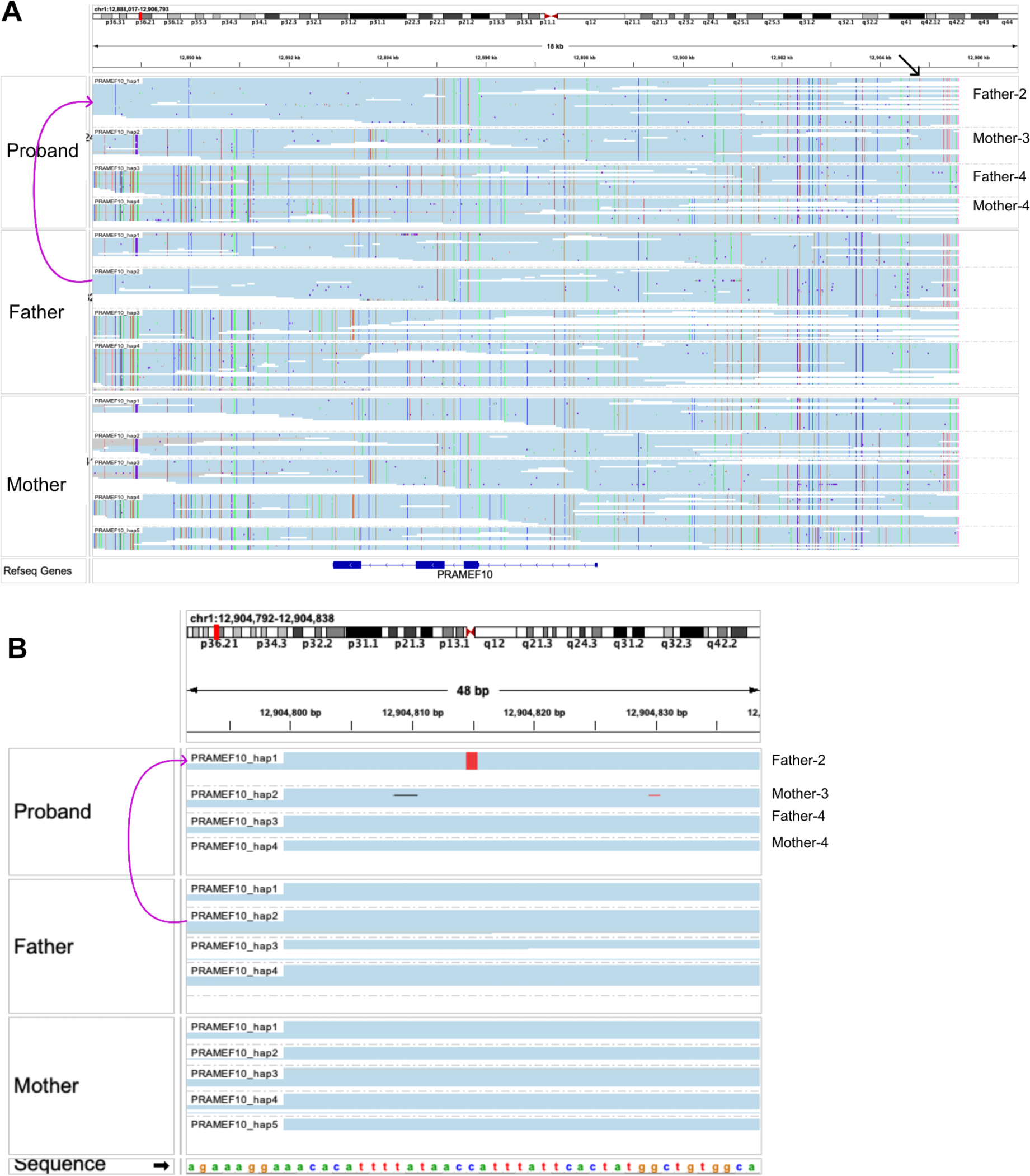
*De novo* SNV #3. Variant is intergenic. Variant is marked by the black arrow. B is a close view of the variant.

**Figure S9.**
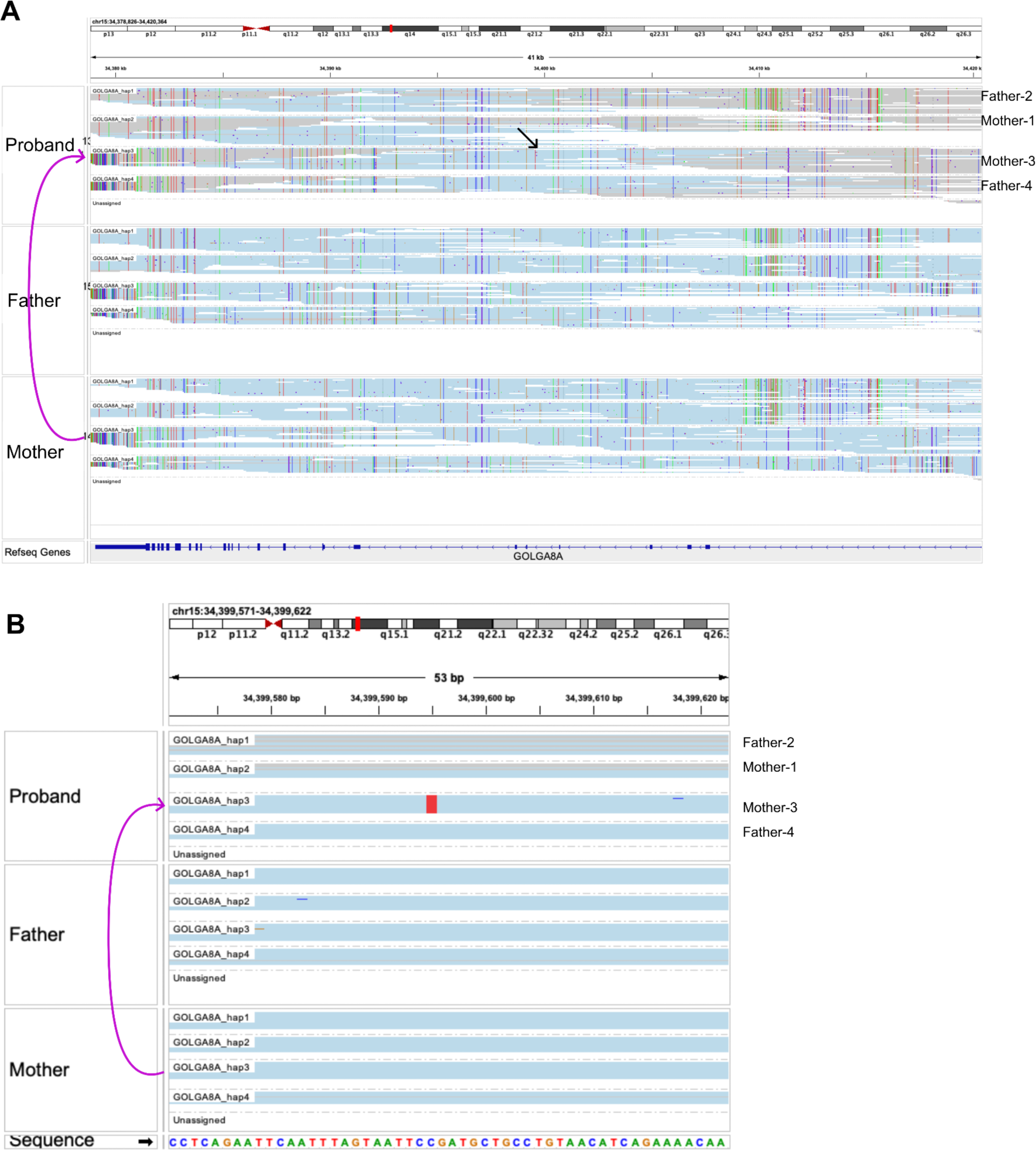
*De novo* SNV #4. Variant is intronic. Variant is marked by the black arrow. B is a close view of the variant.

**Figure S10.**
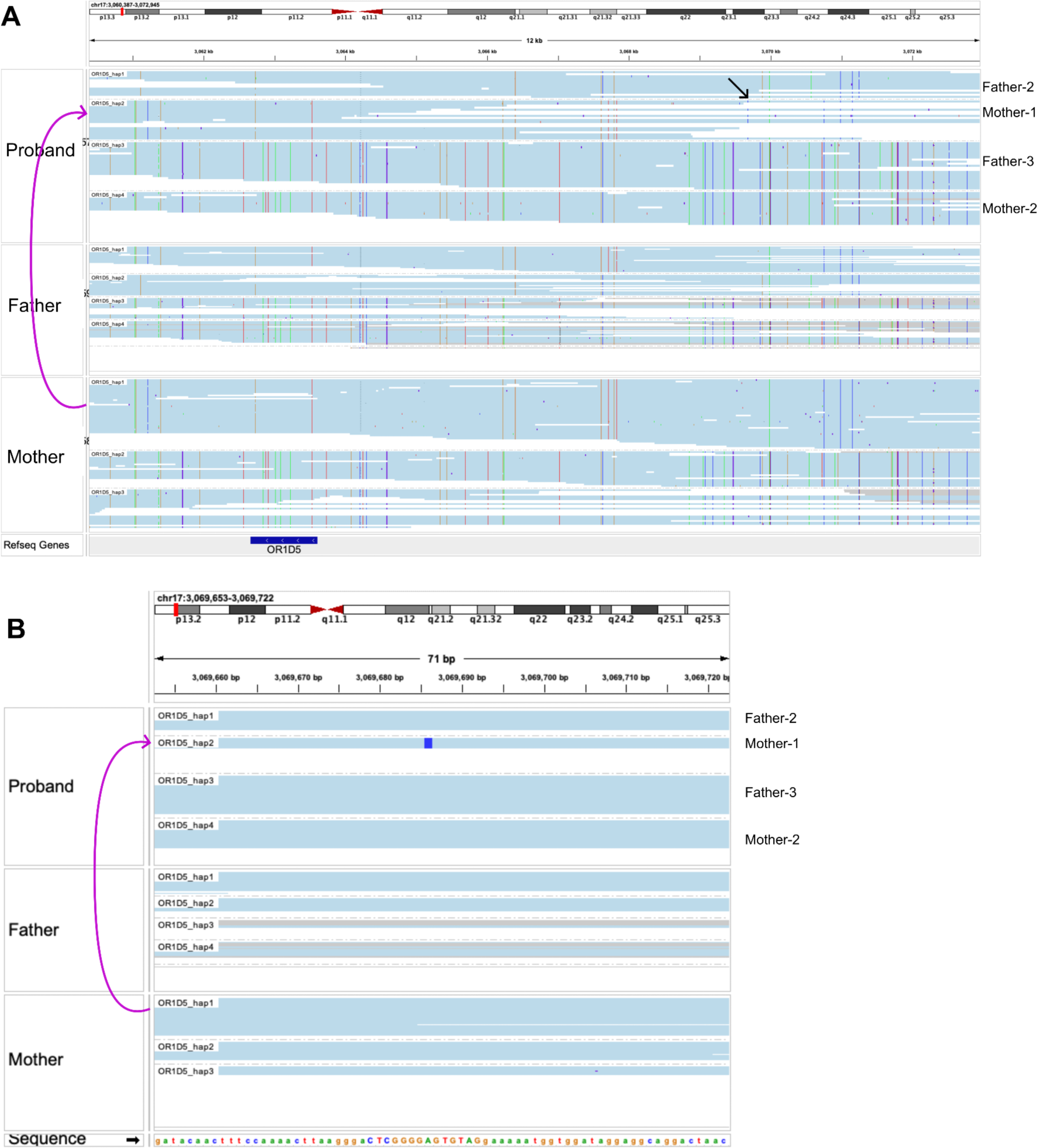
*De novo* SNV #5. Variant is intergenic. Variant is marked by the black arrow. B is a close view of the variant.

**Figure S11.**
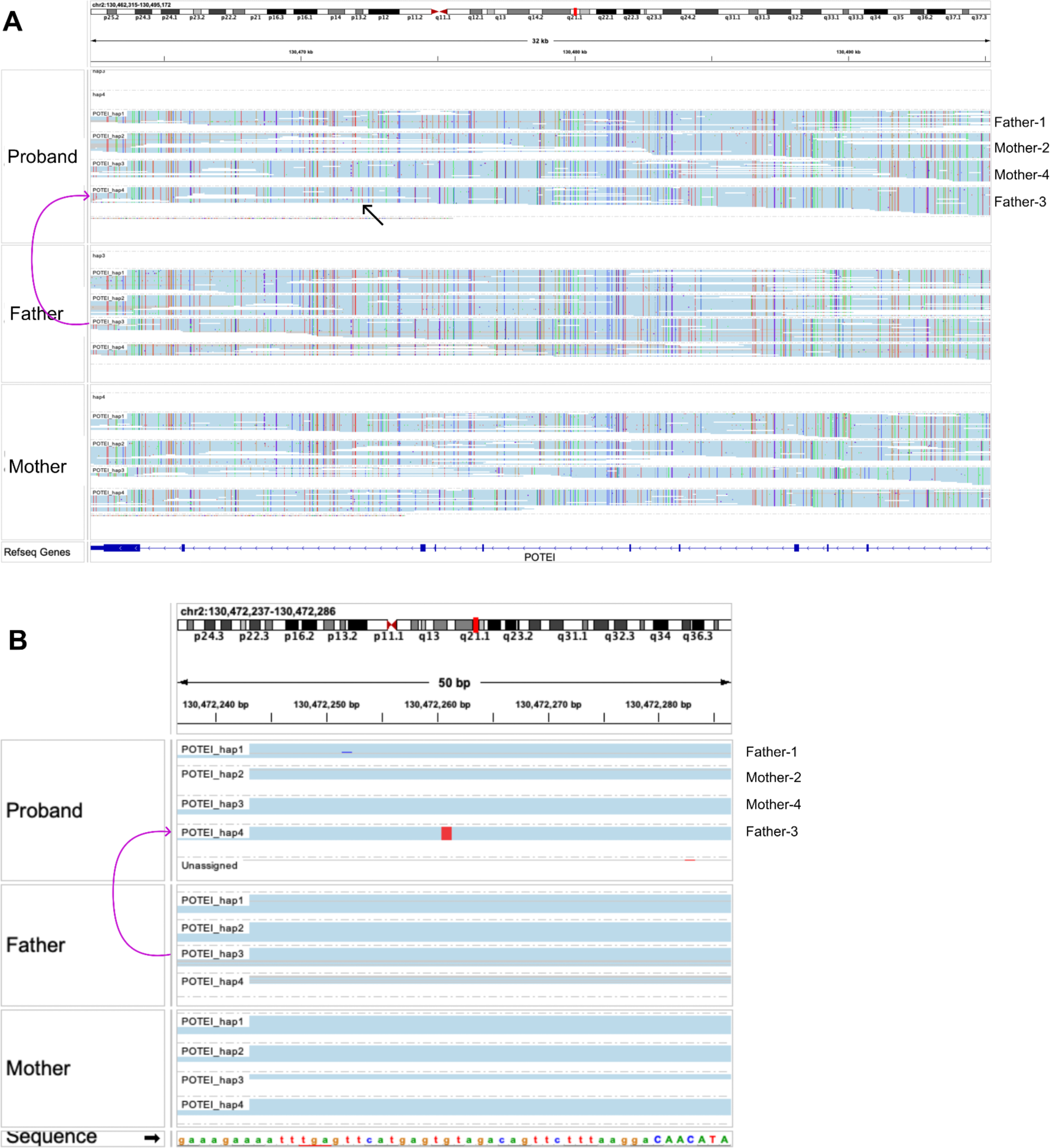
*De novo* SNV #6. Variant is intronic. Variant is marked by the black arrow. B is a close view of the variant.

**Figure S12.**
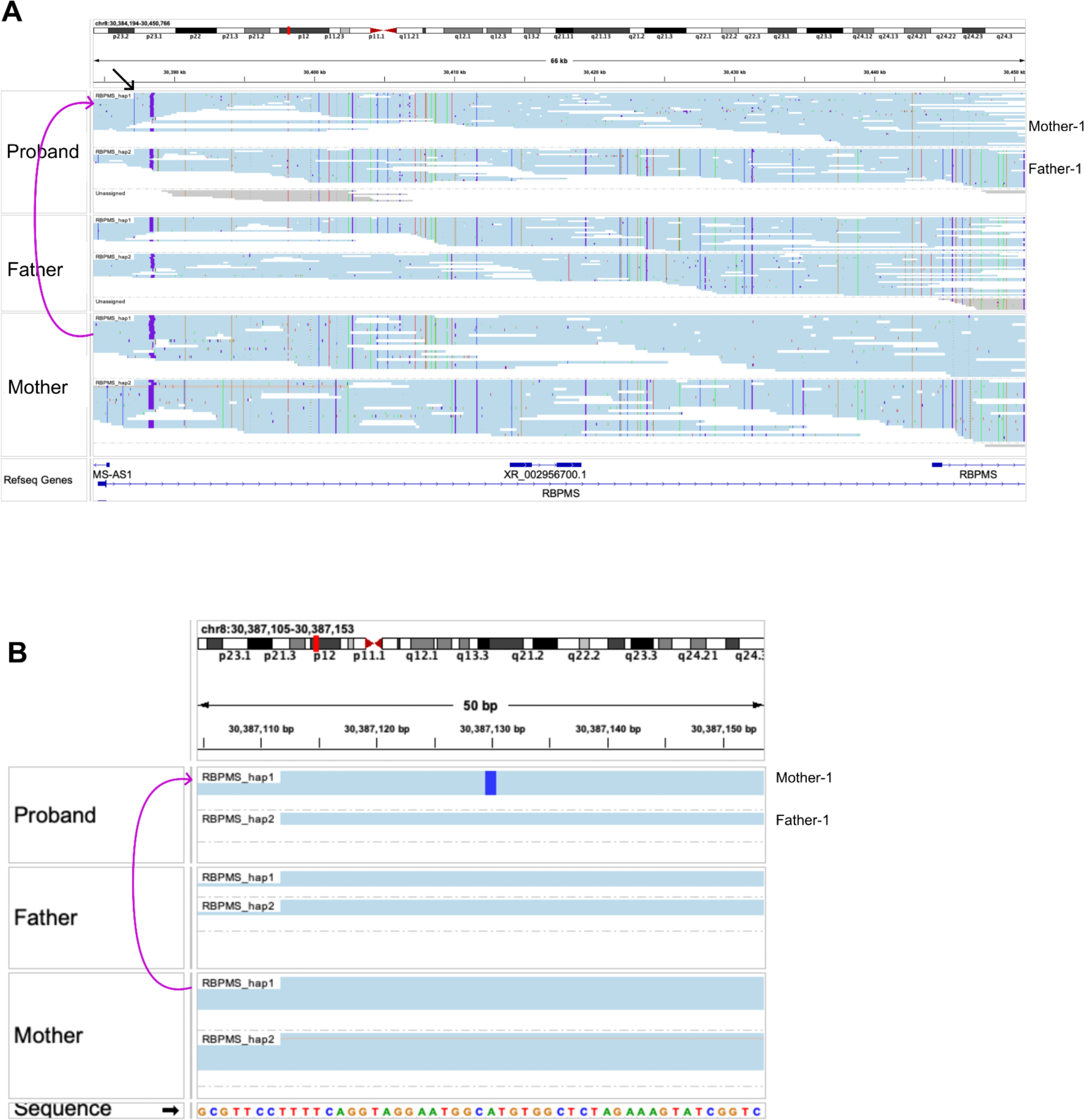
*De novo* SNV #7. Variant is intronic. Variant is marked by the black arrow. B is a close view of the variant.

**Figure S13.**
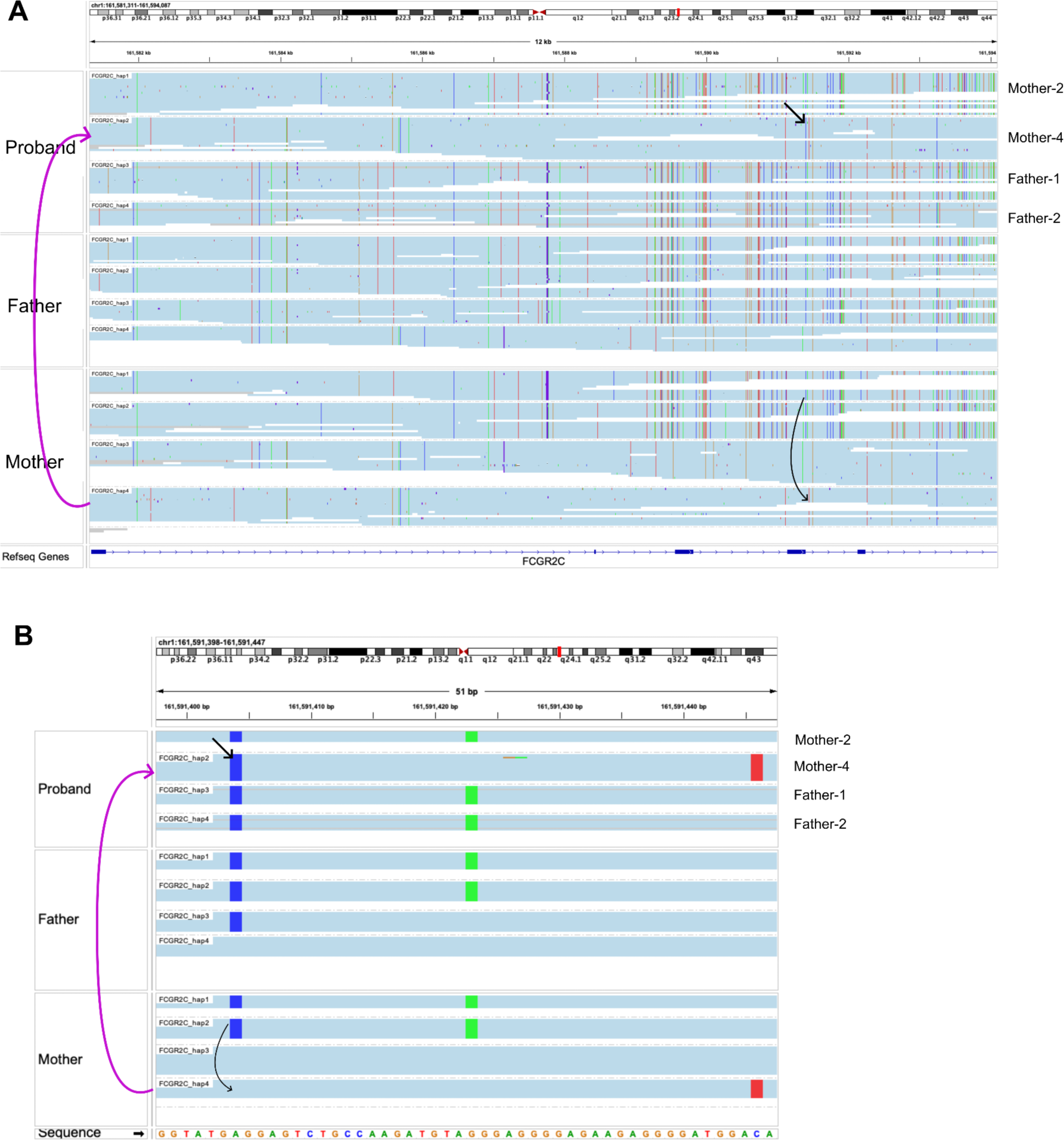
Non-allelic gene conversion #2 (in addition to Figure 4). Variant is intronic. Black arrows mark the variant as well as the potential direction of conversion. B is a close view of the variant.

**Figure S14.**
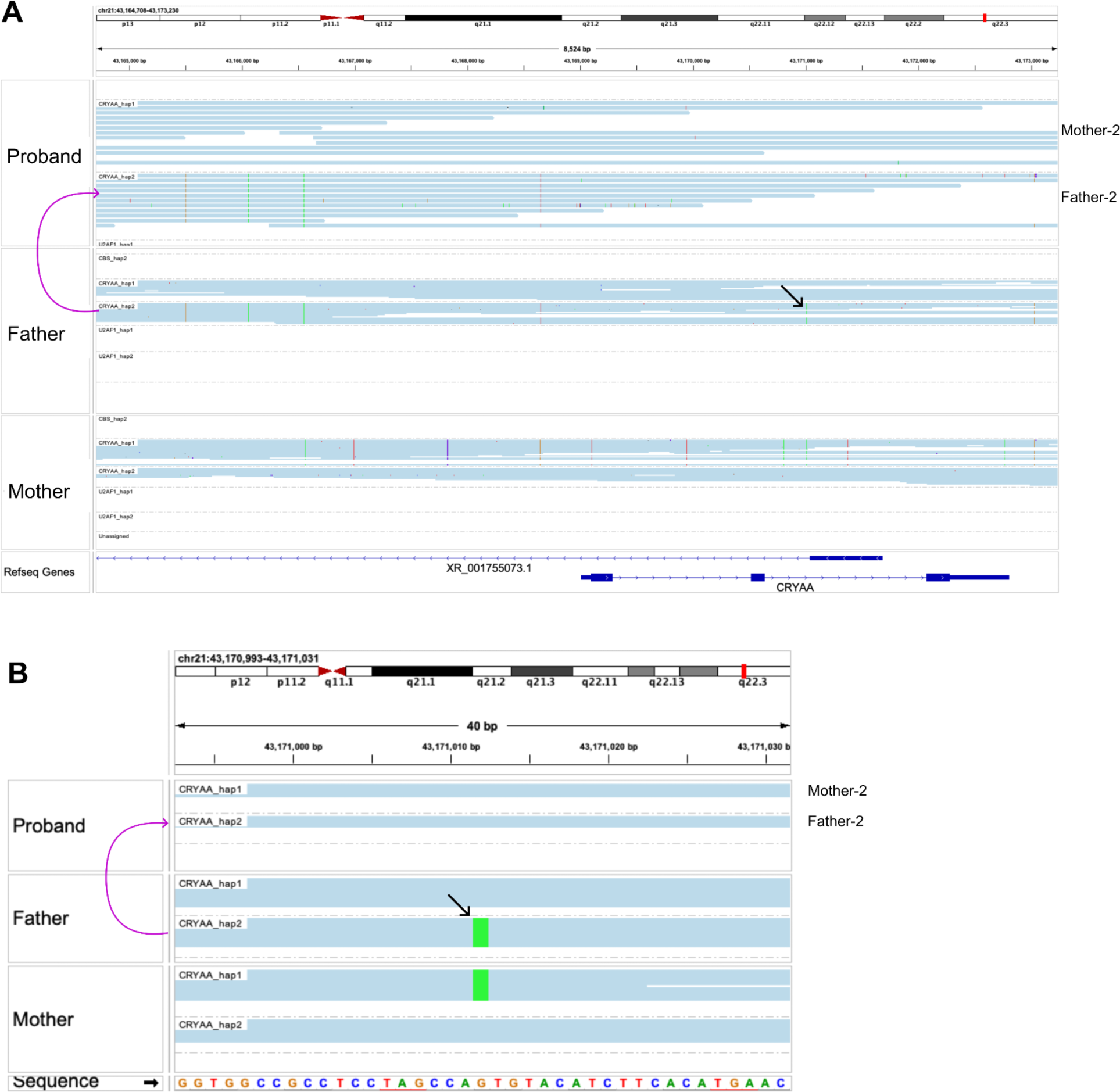
Allelic gene conversion. Variant is intronic. Variant is marked by the black arrow. B is a close view of the variant.

**Figure S15.**
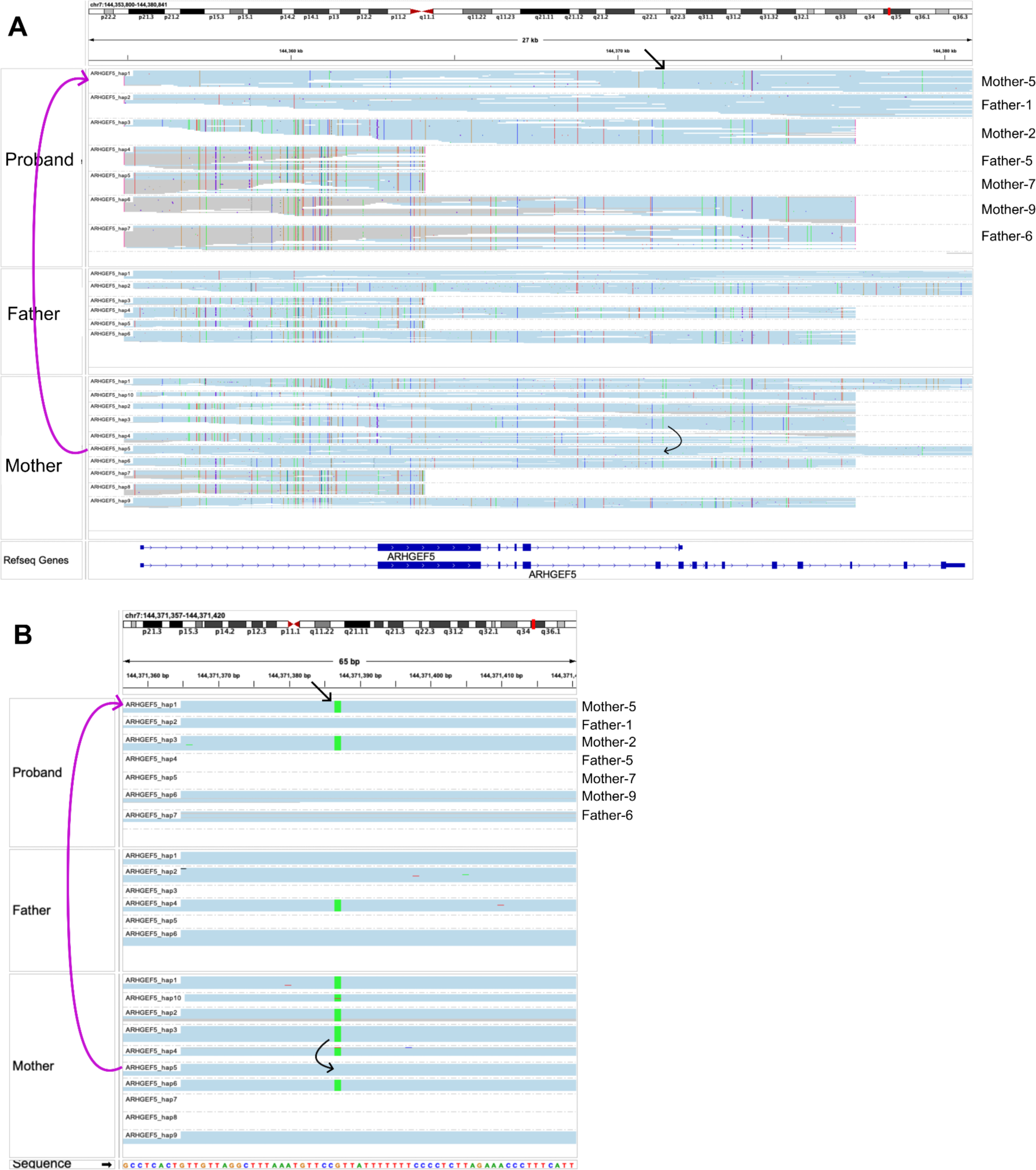
Gene conversion allelic/non-allelic. Variant is intronic. Black arrows mark the variant as well as the potential direction of conversion. B is a close view of the variant.

**Figure S16.**
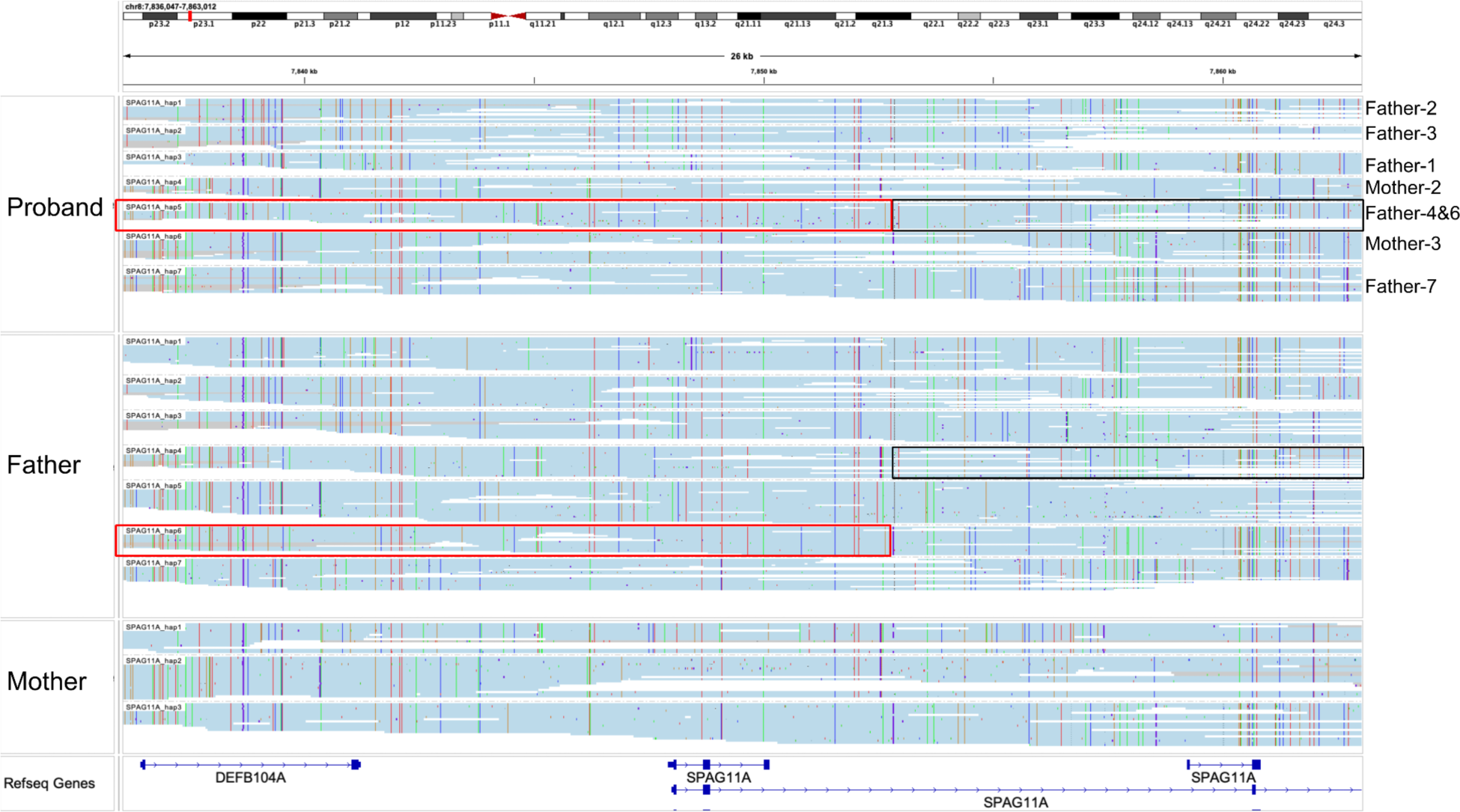
Hybrid haplotype due to equal or unequal crossing over. The red and black boxes mark two parts of the hybrid haplotype and their origin in the father.

**Figure S17.**
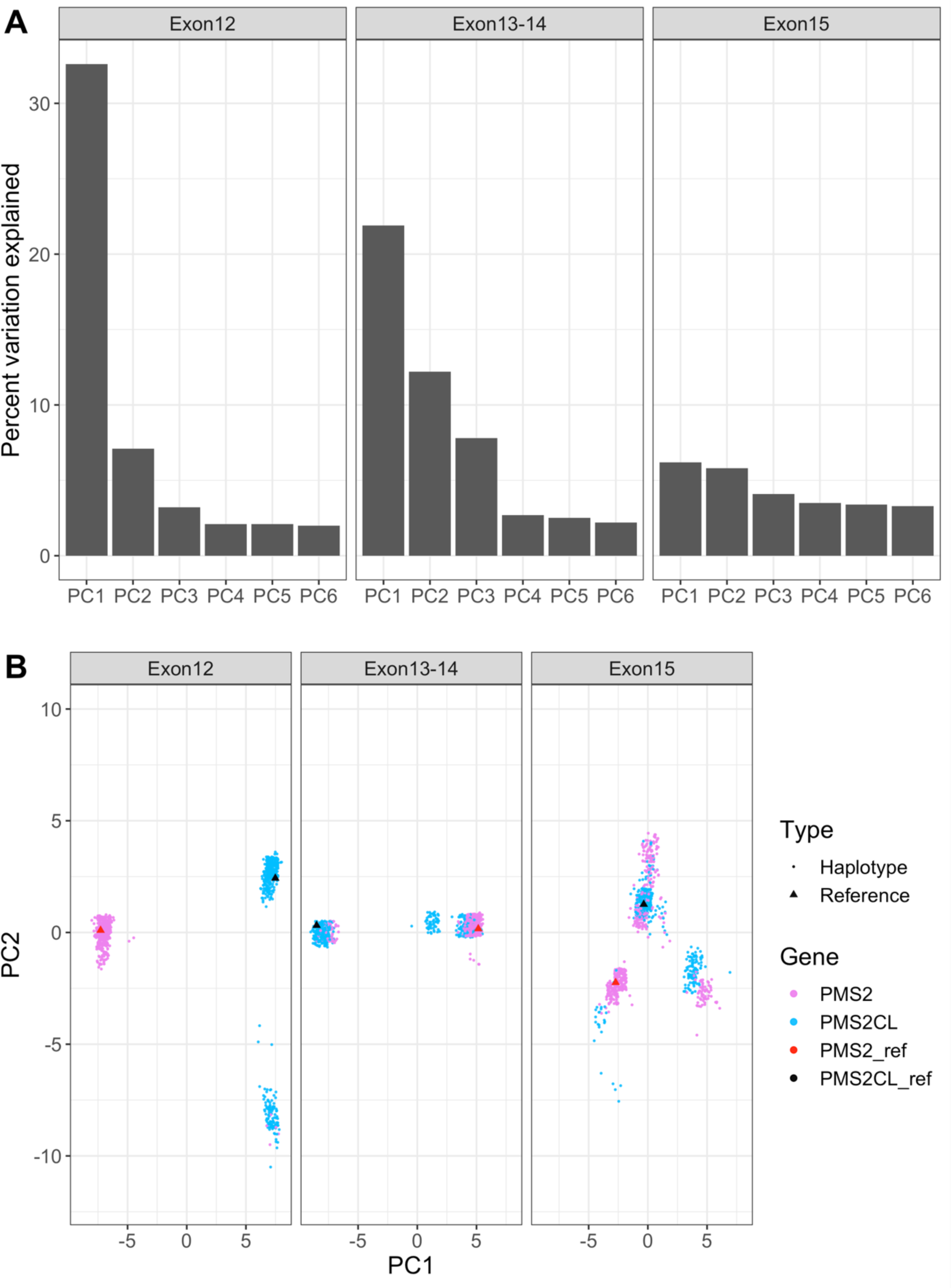
PCA of Exon12, Exon13-14 and Exon15 sequences in *PMS2/PMS2CL*. *PMS2* and *PMS2CL* are distinguishable in Exon 12 and Exons 13-14, with occasional gene conversions. *PMS2* and *PMS2CL* are indistinguishable in Exon 15.

**Figure S18.**
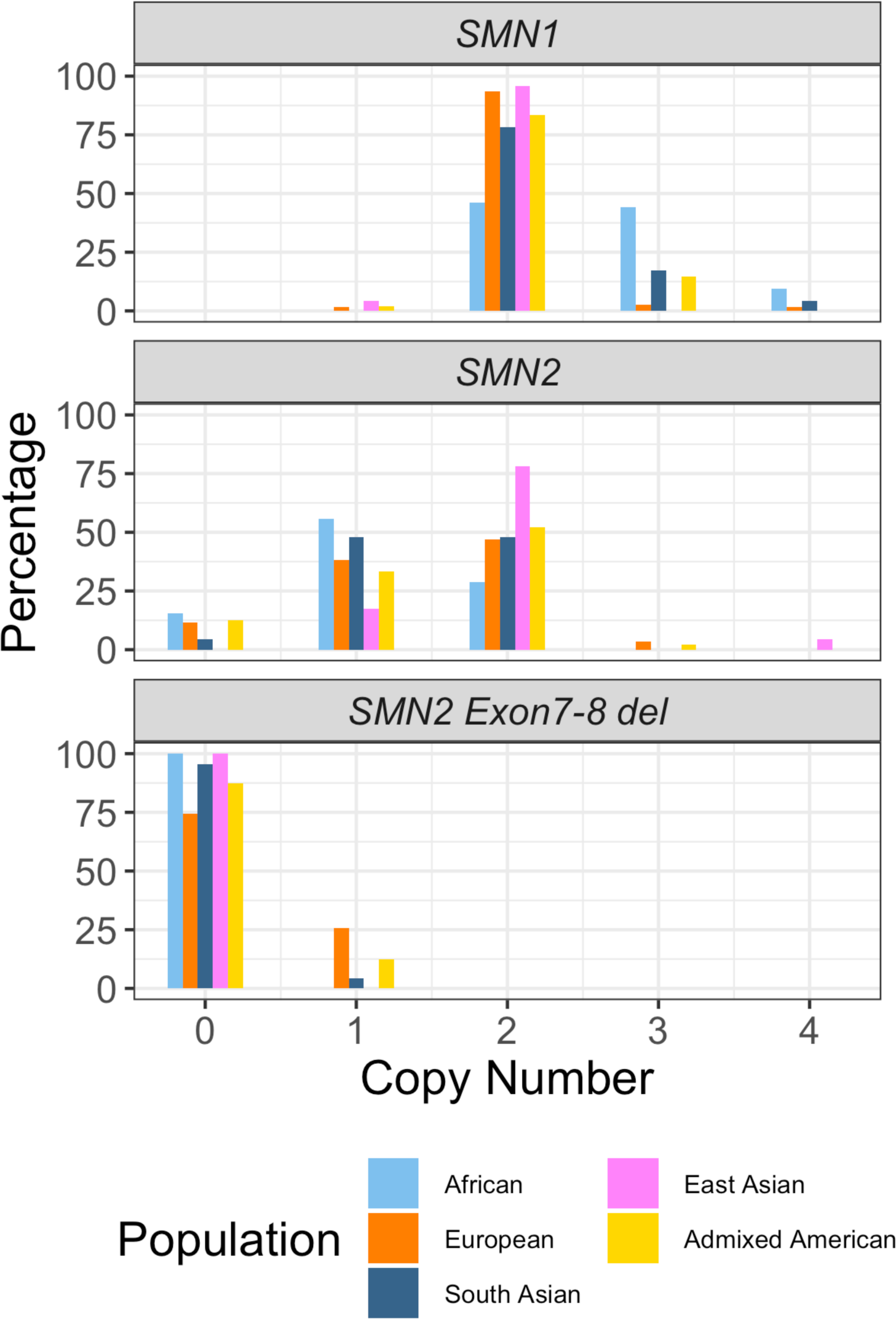
Distribution of the copy number of *SMN1* and *SMN2* across populations.

**Figure S19.**
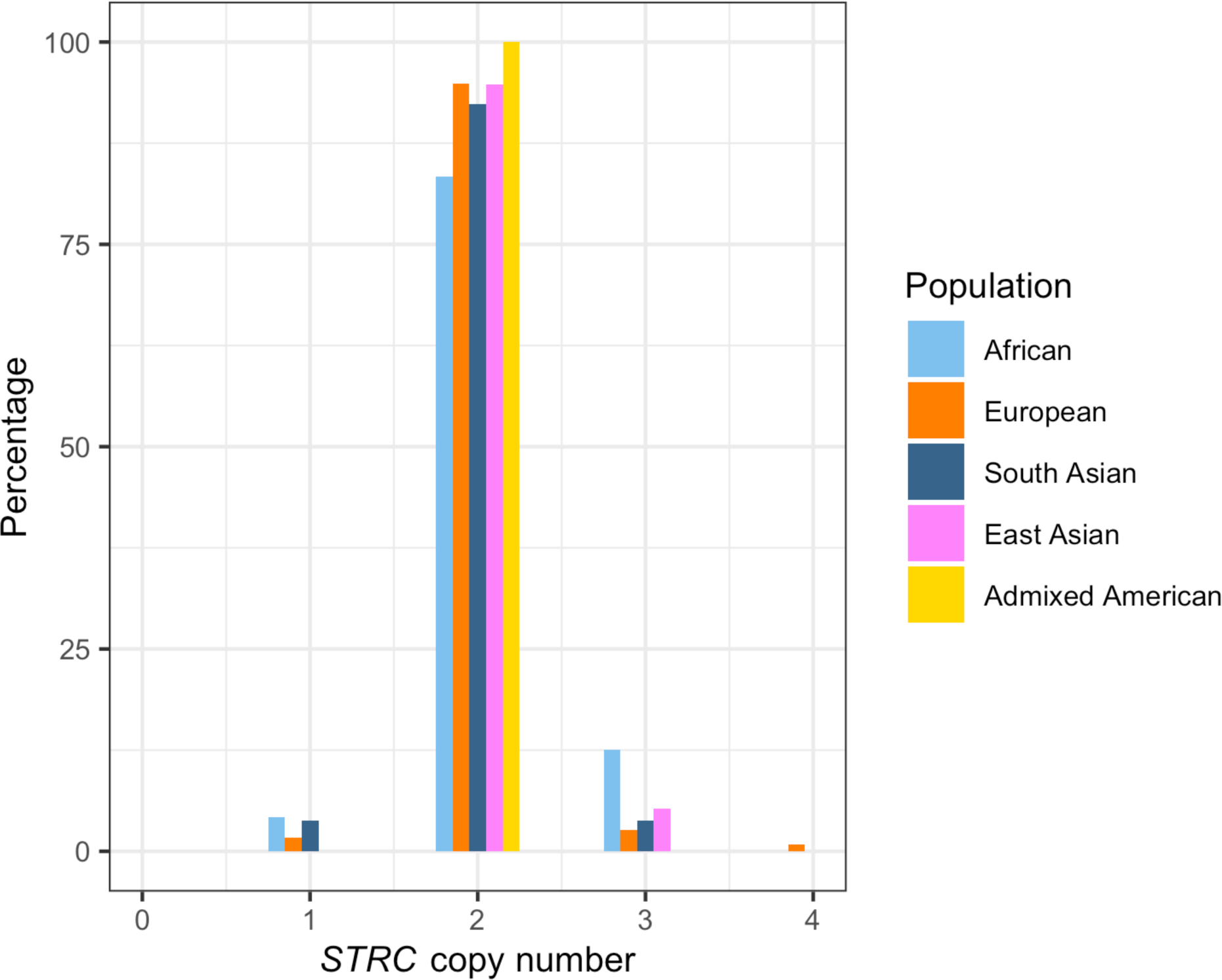
Distribution of the copy number of *STRC* across populations.

**Figure S20.**
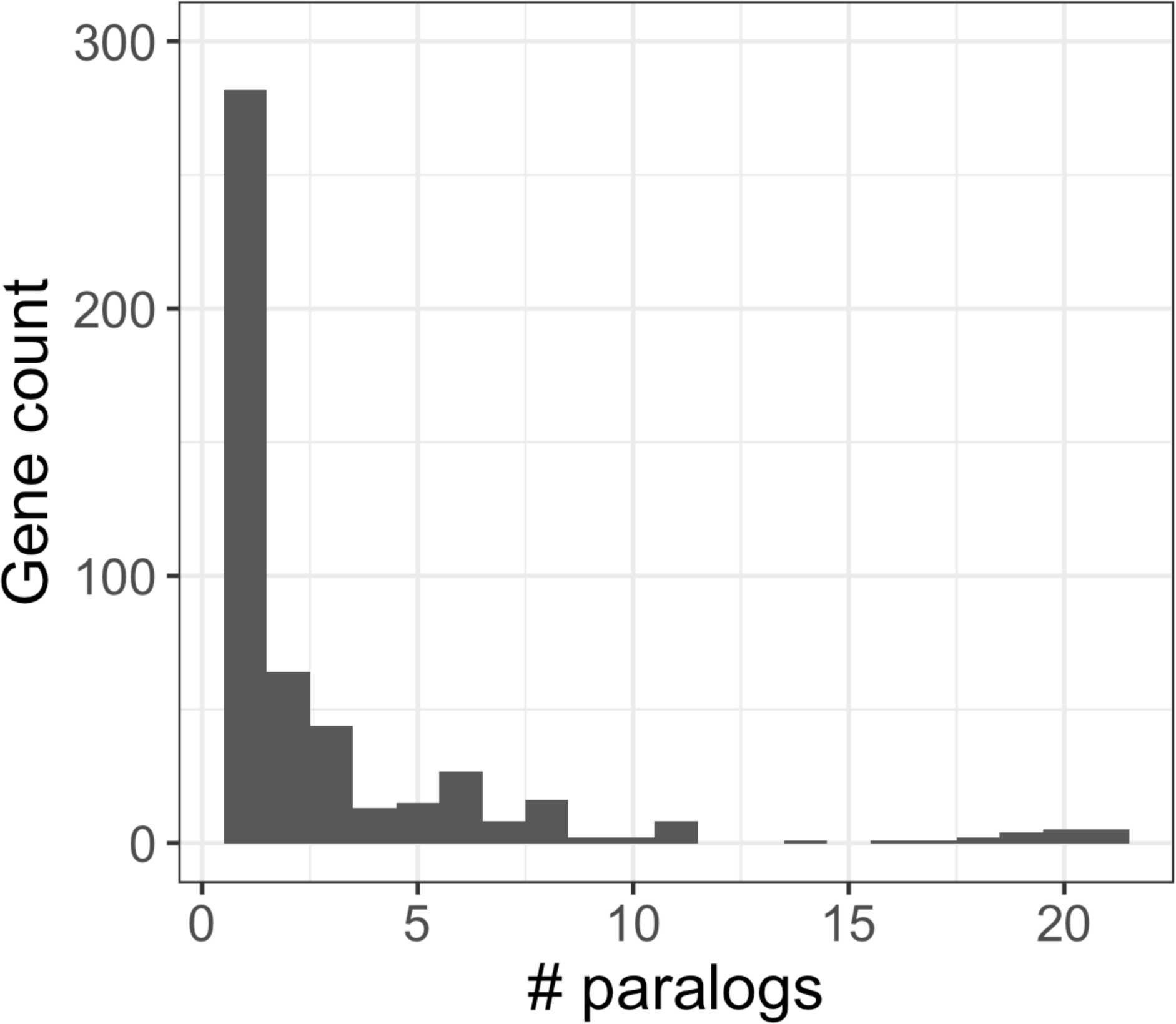
Distribution of the number of highly similar paralogs for genes in SDs.

**Figure S21.**
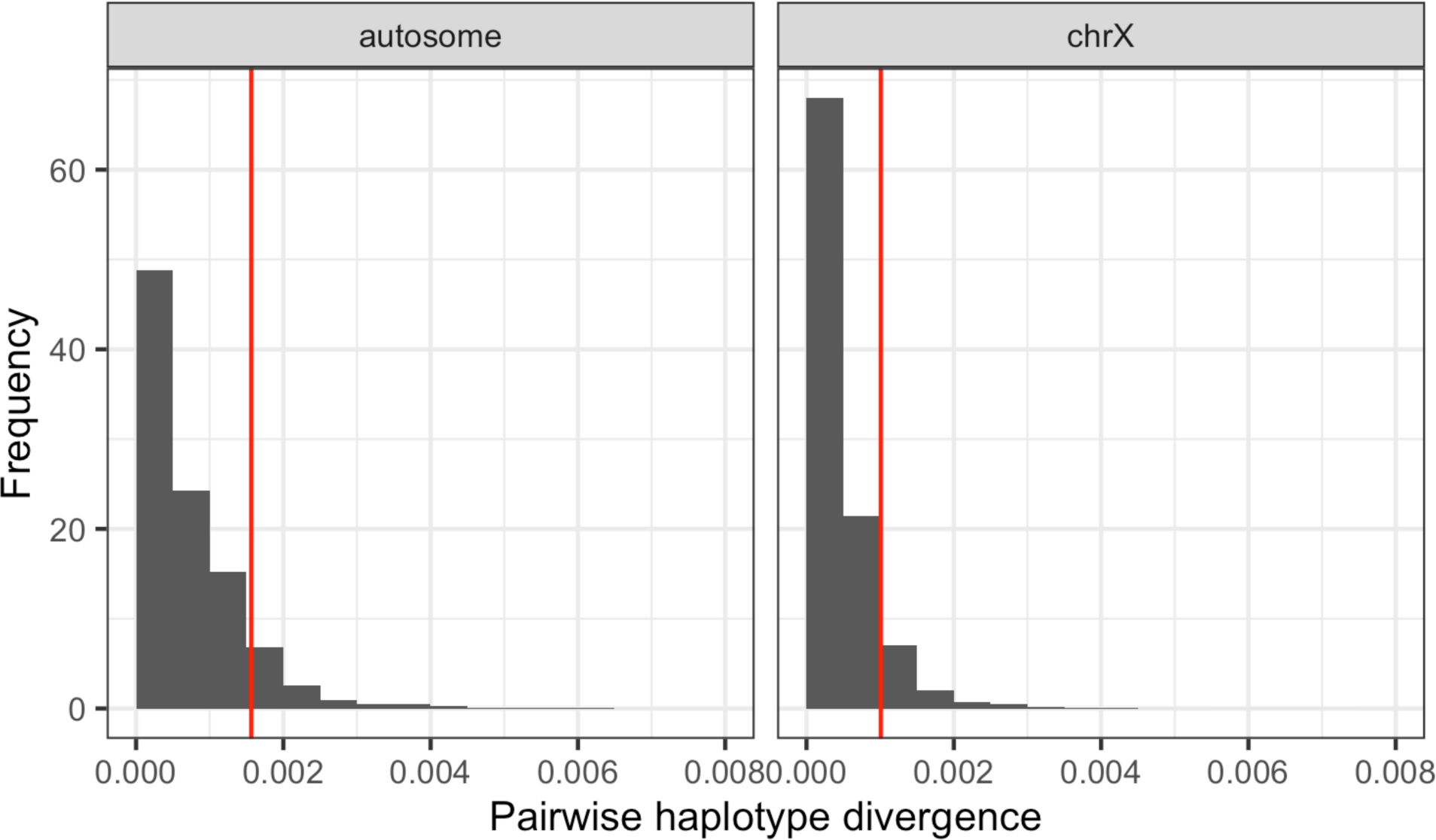
Distributions of allelic sequence divergence for genes without paralogs. Distribution of pairwise sequence divergence between haplotypes among 400 randomly selected autosomal genes and 200 randomly selected chrX genes. Red vertical lines represent the 90th percentile.

### Supplementary Tables

Table S1. Details on SD regions analyzed by Paraphase, including summaries on CN variability and pairwise haplotype divergence (Excel spreadsheet)

Table S2. Details on validation samples (Excel spreadsheet)

**Table S3.**
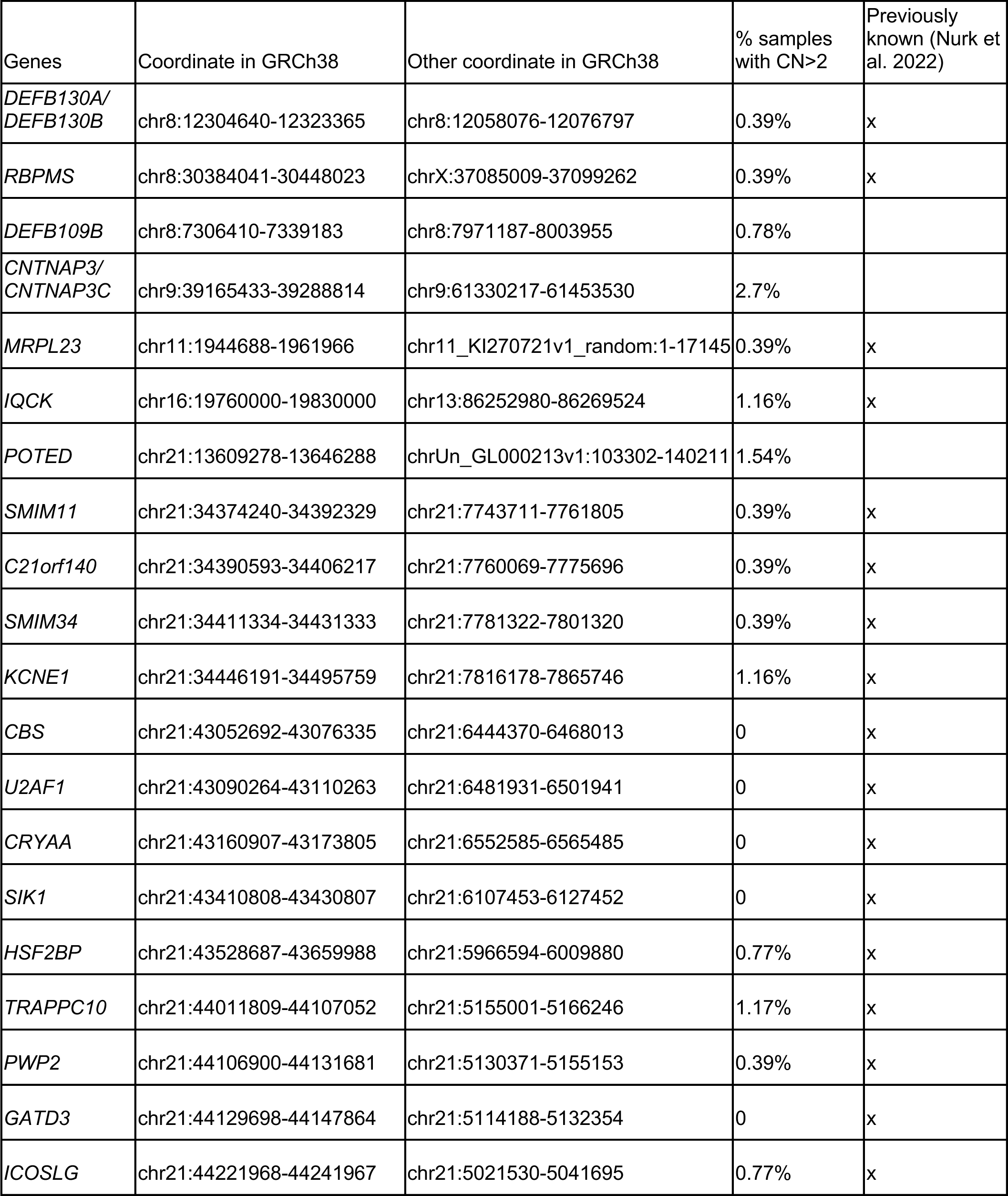
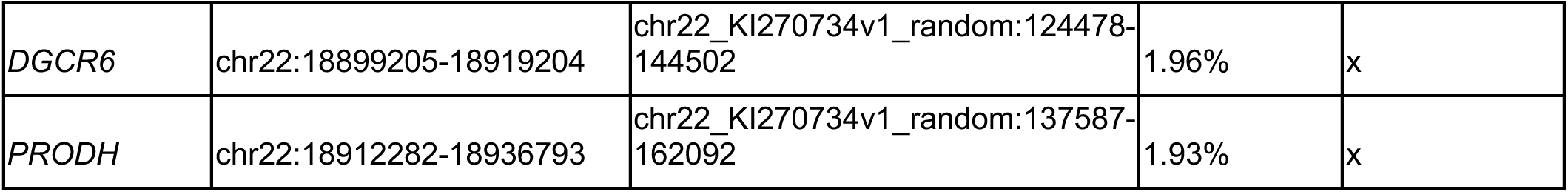
Genes that fall in SDs in GRCh38 but have a consistent CN of two across populations.

**Table S4.** *CYP21A2* allele frequencies.

| Allele | African | European | South Asian | East Asian | Admixed American |
| --- | --- | --- | --- | --- | --- |
| WT (*CYP21A2*, *CYP21A1P*) | 80.2% | 73.5% | 74.0% | 84.2% | 78.9% |
| *CYP21A1P* deletion | 11.5% | 15.0% | 6.0% | 7.9% | 13.2% |
| *CYP21A1P* duplication | 3.1% | 6.6% | 16.0% | 5.3% | 6.6% |
| *CYP21A2* duplication | 1.0% |  |  |  |  |
| *CYP21A2*(WT),<br>*CYP21A2*(Q319X), *CYP21A1P* | 1.0% | 2.2% | 2.0% |  | 1.3% |
| Alleles carrying pathogenic<br>*CYP21A2* variants, listed below | 3.1% | 2.5% | 2.0% | 2.6% |  |
| M240K | 1.0% |  |  |  |  |
| Q319X |  | 0.4% | 2.0% |  |  |
| V282L | 2.1% | 1.3% |  |  |  |
| G111Vfs |  | 0.4% |  |  |  |
| Fusion deletion |  | 0.4% |  |  |  |
| Large conversion |  |  |  | 2.6% |  |

**Table S5.** *OPN1LW/OPN1MW* allele frequencies.

| **Allele** (first two copies in the array) | **African** | **European** | **South Asian** | **East Asian** | **Admixed American** | **Phenotype** |
| --- | --- | --- | --- | --- | --- | --- |
| *OPN1LW+OPN1MW* | 96.2% | 94.0% | 96.3% | 95.2% | 93.9% | Normal |
| *OPN1LW* | 1.3% |  |  | 4.8% | 1.5% | Deutan |
| *OPN1LW+OPN1LW* | 2.5% | 3.4% | 3.7% |  | 1.5% |  |
| *OPN1MW* |  |  |  |  | 1.5% | Protan |
| *OPN1MW+OPN1MW* |  | 2.6% |  |  | 1.5% |  |

**Table S6.**
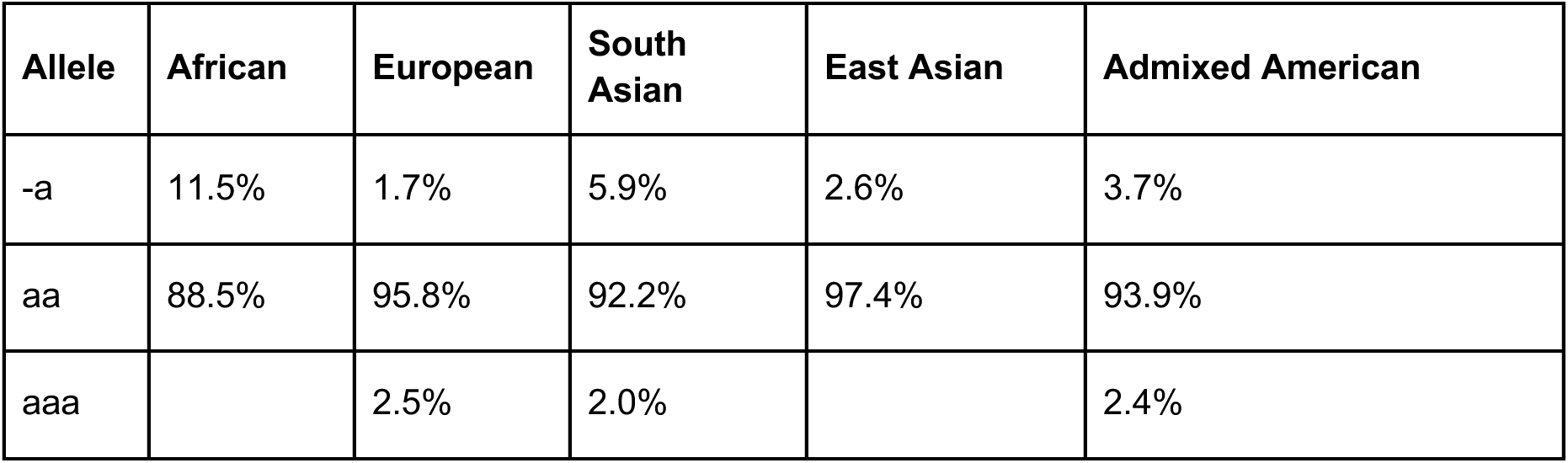
*HBA1/HBA2* allele frequencies.

**Table S7.** Frequencies of the deletion/duplication of the 11.7-kb region (Exons 4-10) in *IKBKG/IKBKGP1*.

| Gene | Allele | African | European | South Asian | East Asian | Admixed American |
| --- | --- | --- | --- | --- | --- | --- |
| *IKBKG* | 11.7-kb deletion |  |  |  |  |  |
|  | 11.7-kb duplication |  | 0.6% |  |  |  |
| *IKBKGP1* | 11.7-kb deletion |  | 0.6% |  |  | 3.2% |
|  | 11.7-kb duplication | 5.1% | 1.2% | 2.8% |  |  |

**Table S8.**
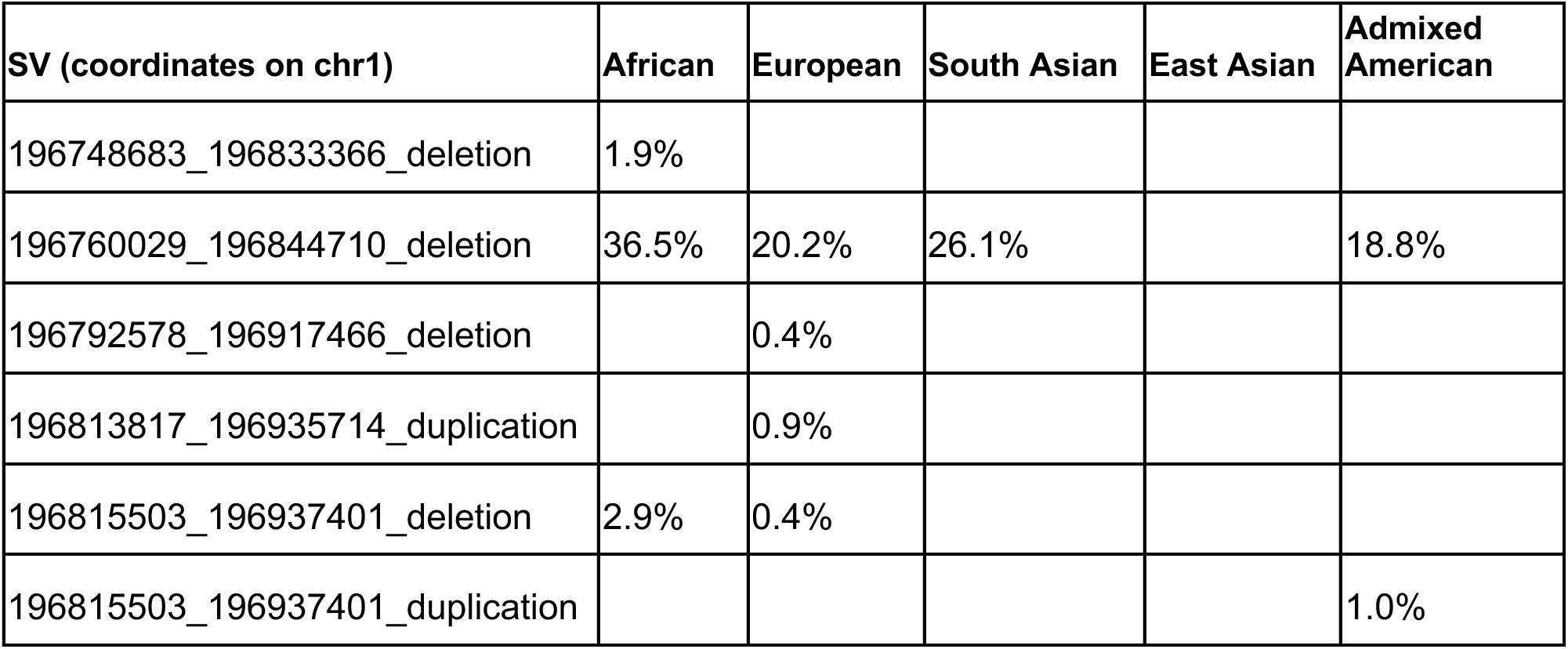
Deletion/Duplication frequencies in the *CFH* gene cluster.

**Table S9.** Deletion/Duplication frequencies in the *GBA/GBAP1* region.

| SV (coordinates on chr1) | African | European | South Asian | East Asian | Admixed American |
| --- | --- | --- | --- | --- | --- |
| 155212923_155233549_deletion |  | 0.4% |  |  |  |
| 155214276_155234903_duplication | 5.8% |  |  |  |  |
| 155215100_155235727_duplication |  |  | 2.2% |  | 1.0% |

## Notes

### Summary of Updates

This version of the manuscript has been revised to update the author list (UCI GREGoR).

